# Genome assembly and analysis of the flavonoid and phenylpropanoid biosynthetic pathways in Fingerroot ginger (*Boesenbergia rotunda*)

**DOI:** 10.1101/2022.05.11.491478

**Authors:** Sima Taheri, Teo Chee How, John S. Heslop-Harrison, Trude Schwarzacher, Tan Yew Seong, Wee Wei Yee, Norzulaani Khalid, Manosh Kumar Biswas, Naresh V R Mutha, Yusmin Mohd-Yusuf, Han Ming Gan, Jennifer Ann Harikrishna

## Abstract

*Boesenbergia rotunda* (Zingiberaceae), is a high-value culinary and ethno-medicinal plant of Southeast Asia. The rhizomes of this herb have high flavanone and chalcone content. Here we report genome analysis of *B. rotunda* together with a complete genome sequence as a hybrid assembly. *B. rotunda* has an estimated genome size of 2.4 Gb which was assembled as 27,491 contigs with N50 size of 12.386 Mb. The highly heterozygous genome encodes 71,072 protein-coding genes and has 72% repeat content, with class I TEs occupying ∼67% of the assembled genome. Fluorescence *In Situ* Hybridization of the 18 chromosome pairs at metaphase showed six sites of 45S rDNA and two sites of 5S rDNA. SSR analysis identified 238,441 gSSRs and 4,604 EST-SSRs with 49 SSR markers common among related species. Genome-wide methylation percentages ranged from 73% CpG, 36% CHG and 34% CHH in leaf to 53% CpG, 18% CHG and 25% CHH in embryogenic callus. Panduratin A biosynthetic unigenes were most highly expressed in watery callus. *B rotunda* has a relatively large genome with high heterozygosity and TE content. This assembly and data (PRJNA71294) comprise a source for further research on the functional genomics of *B. rotunda*, the evolution of the ginger plant family and the potential genetic selection or improvement of gingers.

## Introduction

*Boesenbergia rotunda* (L.) Mansf. (*syn. B. pandurata* (Roxb.) Schltr.) (ITIS Taxonomic Serial No.: 506504), commonly known as Fingerroot ginger and as a type of galanga or galangal, is a member of the family Zingiberaceae in the order Zingiberales. With 50 genera and 1,600 species, the Zingiberaceae is the largest family in the order, along with other families of ginger (Zingiberaceae, Costaceae, Marantaceae, Cannaceae) and banana (Musaceae, Strelitziaceae, Lowiaceae, Heliconiaceae) that include many economically important plant species ^1, 2^. The Zingiberaceae family consists of herbaceous perennial plants that are distributed over tropical and subtropical regions with the highest diversity in Southeast Asia (especially Indonesia, Malaysia and Thailand), India and Southern China ^3–6^. The leaves, flowers and in particular the rhizomes of many of the Zingiberaceae family members are used as flavouring agents and for herbal medicine ^4, 7^.

Boesenbergia is a genus of about 80 species, distributed from India to Southeast Asia ^3, 8–10^. *B. rotunda* is a perennial herb propagated via rhizomes and widely cultivated commercially for its rhizomes and shoots to flavour food and for ethno-medicinal use ^11, 12^. Research on the secondary metabolites of *B. rotunda* has focused on the medicinal properties of rhizome extracts, in particular flavanones and chalcones including panduratin A, pinocembrin, pinostrobin, alpinetin, boesenbergin, cardamonin, naringenin, quercetin, and kaempferol ^8, 13–19^. Of these, the flavonoid compounds panduratin A/DI, 4-hydroxypanduratin, and cardamonin show the clearest biological and pharmacological effects such as anti-inflammatory ^20, 21^, anti-tumor activity against human breast and lung cancers ^22–26^, and antimicrobial activity against HIV protease ^27^, Dengue-2 (DEN-2) virus NS3 protease ^28, 29^, SARS-CoV-2 in human airway epithelial cells ^30^, the oral bacteria *Streptococcus mutans* ^31, 32^, *Helicobacter pylori* ^33^, and against the spoilage bacteria *Lactobacillus* (*Lactiplantibacillus*) *plantarum* ^34^. A recent patent claimed that panduratin derivatives from *B. rotunda* have potential for preventing, ameliorating, or treating bone loss disease ^35^, while 4-hydroxypanduratin was reported to have the most potent vasorelaxant activity among the major flavonoids of *B. rotunda* extracts ^36^.

The ethnomedicinal and potential pharmaceutical importance of *B. rotunda* have led to interest in exploring cell and tissue culture for secondary metabolite production. In commercial farms, the plant is propagated clonally from rhizomes, and several protocols for multiplication via *in vitro* culture have been reported including plantlet regeneration *via* somatic embryogenesis from callus cultures ^37, 38^, from shoot bud explants ^39^ and from embryogenic cell suspension cultures ^38^. Cell suspensions of *B. rotunda* ^17, 40^ and various types of callus ^41, 42^ have been explored as potential sources for alpinetin, cardamonin, pinocembrin, pinostrobin and panduratin A. Reproducible methods for *in vitro* cell culture of *B. rotunda*, led to protocols for genetic transformation ^38^, that could facilitate metabolic engineering of cell materials for specific desirable metabolite production. However, current knowledge of the underlying biosynthetic pathways is sparse. Other than biochemical profiling ^16, 42^, the application of current technologies for determining deep sets of the genetic sequences expressed in various tissue and cell types can deliver useful information.

Genomic level studies improve understanding of the biology and biochemistry of the plant and can be applied in breeding for improved agronomy and plant products. Whole genome sequencing identifies genes and regulatory sequences for complex biological processes such as secondary metabolite biosynthesis ^43–45^, while transcriptional profiling provides information for functional studies. Structural genomic studies have been undertaken in other Zingiberales including turmeric (*Curcuma longa*; genome size of 1.24 Gb) ^46^ and for several Musaceae species and cultivars, which have genome sizes ranging from 462 Mb to 598 Mb (Banana Genome Hub https://banana-genome-hub.southgreen.fr/)^47^; while the Pan-genome of Musa Ensete has a genome size of 951.6 Mb ^48^. Larger plant genomes have now been sequenced including those of important monocot species such as wheat ∼ 17 Gb (International Wheat Genome Sequencing Consortium), *Aegilops tauschii* ∼ 4.3 Gb ^49^, oil palm ∼ 1.8 Gb ^50^ and maize ∼ 2.6 Gb ^51^, in addition to species known for their unique metabolites such as tea *(Camellia sinensis)* ∼ 2.98 Gb ^52, 53^ and ginseng (*Panax ginseng*) ∼ 3.2 Gb ^54^. However, even with the recent advances in long sequence technology, large plant genomes can be challenging to assemble due to high repeat content and high levels of heterozygosity ^55, 56^.

The availability of an assembled genome sequence expands the functional biological questions that can be asked, since regulatory and variable elements, many of which may be involved in epigenetic regulation, cannot be seen purely using expression data. So while transcriptome ^40^ and proteome ^57^ data for *B. rotunda* are available, the lack of a previously published genome assembly is a limitation for functional studies. Genome assemblies also facilitate the exploration of genomic repeats which can not only be a source for genetic markers but are also drivers of genome size, gene content and order, centromere function and reflect genome evolution ^58, 59^. Last but not least, the epigenetic dynamism in genomes mainly involves “non-coding DNA” thus a genome assembly provides the framework for epigenetic studies. Therefore, in the current investigation we performed the first complete genome sequence for *B. rotunda* made with a hybrid assembly strategy using Pacific Biosciences (PacBio) and Illumina HiSeq platforms. We explored the sites of 45S rDNA and 5S rDNA on metaphase chromosomes observed by Fluorescence *In Situ* Hybridization (FISH). In addition, we carried out a deep transcriptome (RNA-seq data) assembly from five *B. rotunda* samples, including various types of callus cultures, and leaves. Gene expression profiles and bisulfite seq DNA methylation data from these tissues and samples were used for co-expression analysis to identify any association of gene expression and local DNA methylation of unigenes related to methylation, somatic embryogenesis, and pathways for flavonoid and phenylpropanoid biosynthesis. We also report novel expressed sequence tags-SSR (EST-SSR) and genomic SSR markers for *B. rotunda* and the estimated cross-transferability of the designed primers between *B. rotunda* and closely related species to provide deeper genetic resources to support further study of the biology and biodiversity in this genus. Genomic information and complete sequence data for this less investigated herb should provide a solid foundation as a vital step in genetic analysis to facilitate *B. rotunda* improvement and to reach a deeper understanding of the metabolic pathways of its natural products.

## Results

### Chromosomes and location rDNA sites

*Boesenbergia rotunda* (2n=36; 18 pairs of submetacentric chromosomes) has 3 pairs of 45S rDNA sites near the ends of three pairs of chromosomes (Fig. 1a). One pair of 5S rDNA sites (Fig. 1d) are on a chromosome pair not bearing 45S rDNA.

**Figure 1.**
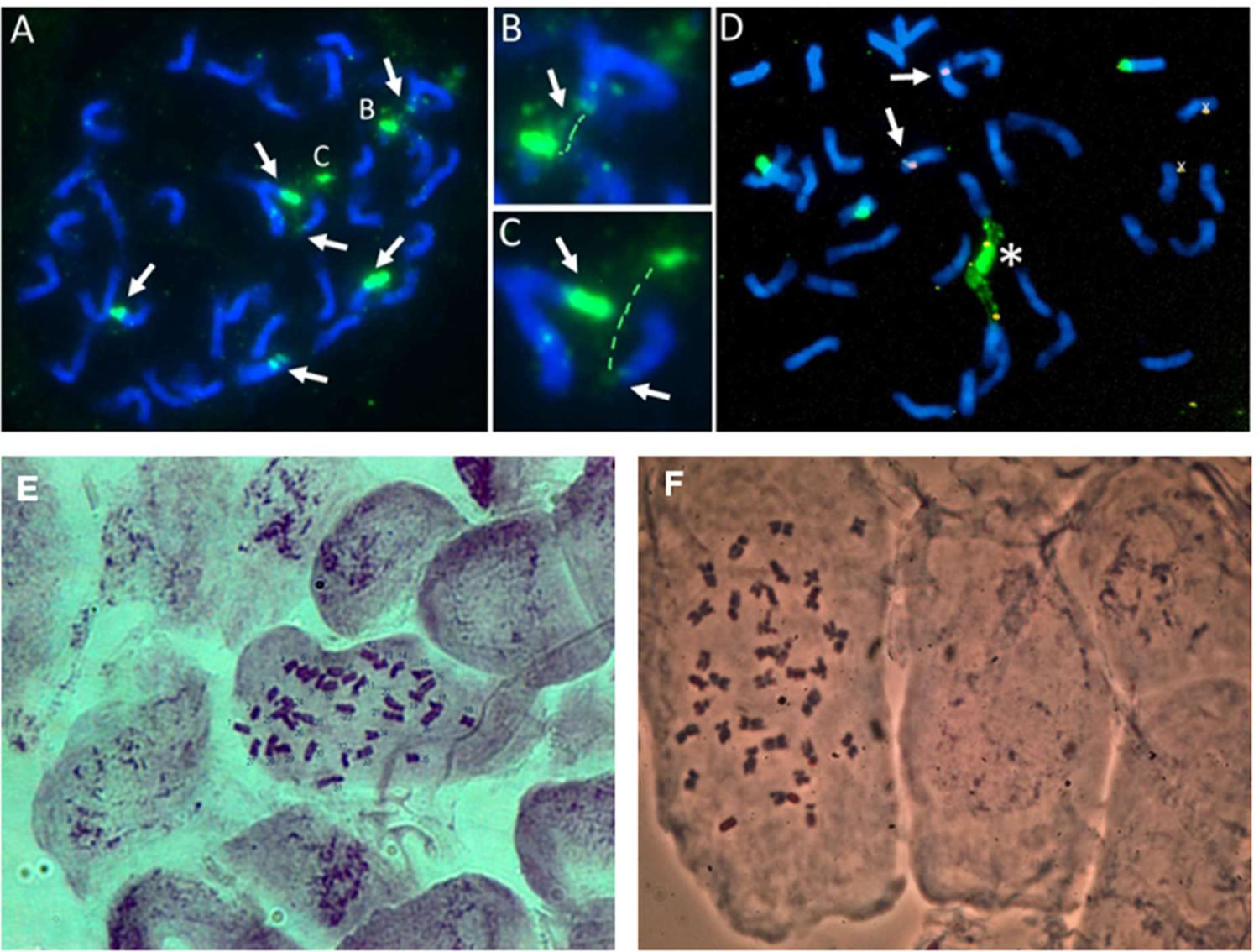
Number and location of 45S and 5S rDNA sites on metaphase chromosomes of *Boesenbergia rotunda (*2n=36). Fluorescent *in situ* hybridization with clone pTa71 (45S rDNA of wheat), labelled with digoxigenin and detected with FITC (green) and clone pTa794 (5S rDNA of wheat) labelled with biotin and detected with Alexa 674 (shown in red). A-C: early metaphase showing 6 sites of 45S rDNA (arrows) of variable strength at ends of 3 pairs of chromosomes. In some cases, the rDNA is extended, and the satellite is separated from the main chromosomes shown enlarged in B and C. D: Two 5S rDNA sites (arrows) were detected on a chromosome pair not bearing 45S rDNA. The star indicates fusion of 2 or 3 45S rDNA sites. E and F: Chromosome preparation using fresh root tips from plants grown and analysed in two different laboratories (E: University of Malaya and F: University of Leicester) showing 36 chromosomes.

### Genome assembly

Genomic DNA from leaves of a single, clonal *B. rotunda* plant was sequenced using multiple approaches (Table S1), with 114 Gb PacBio long reads, 260 Gb of Illumina HiSeq 2500 250bp paired-end reads, and 90 Gb of mate-paired reads with 2, 5, 10, 20 and 40 kb insert sizes. Based on k-mer analysis (k=17, GenomeScope), the estimated haploid genome size of *B. rotunda* was 2.4Gb (Fig. S1), consistent with flow cytometry (Fig. S2). The heterozygosity was estimated as 3.01%. A hybrid genome assembly pipeline combining Illumina data and PacBio data was adopted (Fig. S3). The final assembled genome size was 2.347Gb characterized by 27,491 contigs and 10,627 scaffolds, with contig N50 of 123.86 kb and scaffold N50 of 394.68 kb (Table 1). Based on benchmarking universal single-copy orthologs (BUSCO) analysis ^60^ mapping the *B. rotunda* genome against a set of 1,440 core eukaryotic genes, 1,232 (85.6%) were present (Table S2). Assembly quality assessment showed over 95% of Illumina PE250 reads to map to the contig assembly (Table 2).

**Table 1:** Statistics of the final genome assembly of the *B. rotunda*

|  |  | Scaffolds |  | Contigs |  |  |
| --- | --- | --- | --- | --- | --- | --- |
|  | No. | Size (bp) |  | No. | Size (bp) | Gaps |
|  |  | With gaps | Without gaps |  |  |  |
| Total Number | 10,627 |  |  | 27,491 |  | 16,864 |
| Min | - | 5,830 | 5,830 | - | 5,198 | 25 |
| Median | - | 136,187 | 131,005 | - | 55,047 | 2,415 |
| Mean | - | 220,901 | 213,350 | - | 82,473 | 4,758 |
| Max | - | 2,848,924 | 2,758,809 | - | 1,033,476 | 38,914 |
| Total size | - | 2,347,517,452 | 2,267,274,222 | - | 2,267,274,222 | 80,243,230 |
| N50 | - | 394,682 | 379,106 | - | 123,867 | 11,038 |
| N90 | - | 107,821 | 103,307 | - | 37,045 | 2,551 |
| N95 | - | 69,101 | 66,089 | - | 27,170 | 1,540 |
| GC content (%) | 40.1 |  |  |  |  |  |

**Table 2.** Evaluation of completeness of the final assembly

| Species | Read Length(bp) | Data | Sequence Depth (X) | Mapped (%) | properly paired (%) | singletons (%) | Reference total length (Gb) | Reads covered length (Gb) | Coverage (%) |
| --- | --- | --- | --- | --- | --- | --- | --- | --- | --- |
| *Boesenbergia rotunda* | 250_250 | 260 (Gb) | 104 | 95.24 | 84.47 | 0.20 | 2.35 | 2.25 | 96 |

### Annotation of the *B. rotunda* genome

Five sets of RNA-seq datasets were generated from three cell culture types, *in vitro* and *ex vitro* leaves of *B. rotunda*, given the importance for secondary metabolites production. Individual transcriptomes were assembled from these RNA-seq reads using different *de novo* transcriptome assemblers (Table 3, Fig. S4). The assembled transcriptome size ranged from 31 to 71 million base pairs with 72,085 to 158,465 contigs for the Oases, SOAPdenovo-Trans, TransAbyss, and Trinity (Table 3, Fig. S4). Oases had the highest N50 size and average contig length. The BUSCO quantitative measure of the completeness transcriptomes in terms of expected gene content scores, also showed Oases (36.7%) and TransAbyss (36.6%) to give assemblies with higher numbers of complete and single copy contigs compared to SOAPdenovo-Trans and Trinity (31.7%) (Fig. S5). The non-redundant transcript sequences formed from Oases followed by TGICL were used to annotate the *B. rotunda* genome and for downstream expression analysis.

**Table 3.**
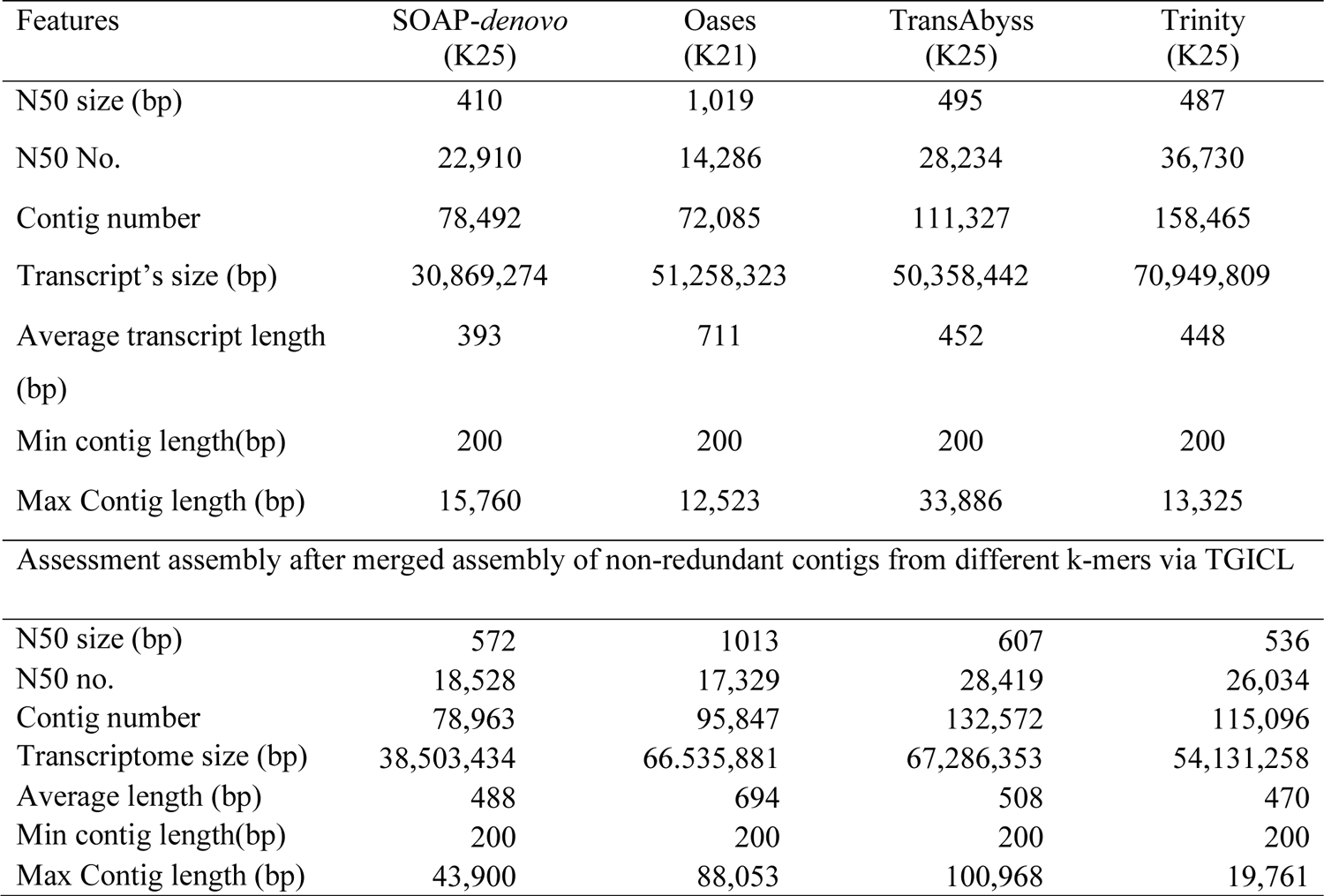
Comparison of *de novo* transcriptome assembly results for four different assembly software: SOAP-*denovo*, Oases, TransAbyss, and Trinity.

Based on a combination of *de novo* and homology-based gene prediction methods, 72.51% of the genome (1.70 Gb) was annotated as repeats including 6.94% tandem repeats. Among Class I TEs (Retroelements), long terminal repeats (LTRs) constituted the greatest proportion of the genome (67.16%) while DNA TE made up 3.29 % of the genome (Fig. S6, Table 4). From 10,627 assembled contigs and 95,847 assembled transcriptome sequences searched for SSRs, (Table 5, Fig. 2), the density of the microsatellites was 102 SSR loci per Mbp in genomic and 69 SSR loci per Mbp in transcriptome sequences. Among the identified repeat motif types, trinucleotides were the most abundant in both genomic (35.62%) and transcriptome (51.67%) sequences, followed by mono- and dinucleotide repeats (Table 5, Fig. 2a). Class II type SSR-loci (<30bp) were two-fold higher than class I type in genomic sequences, whereas class II type SSR-loci were four-fold higher than class I types SSR loci in the transcriptome sequences (Fig. 2b). The number of AT rich microsatellites was significantly higher than that of GC rich and microsatellites with balanced GC content.

**Figure 2.**
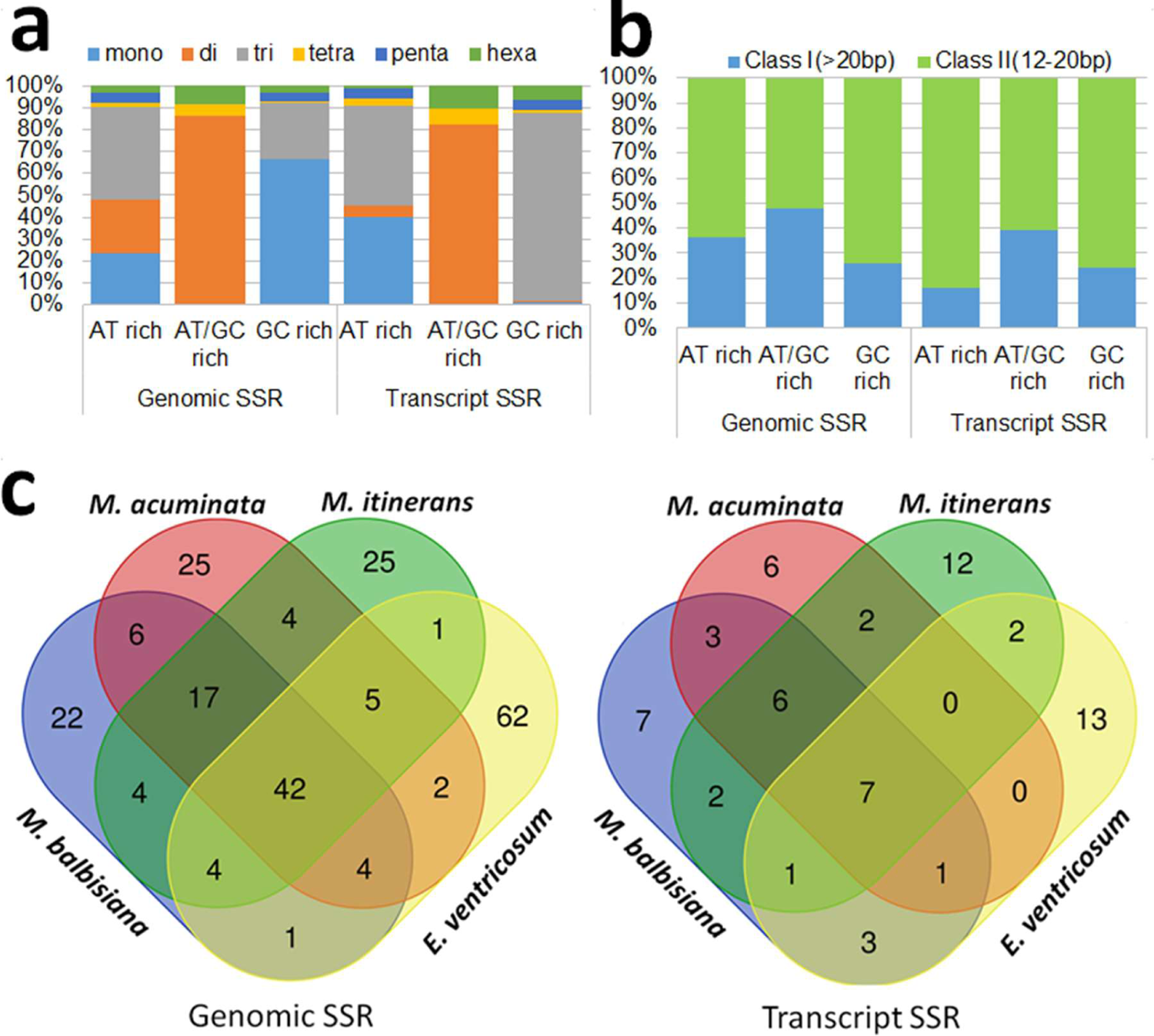
(a-b) Frequency distribution of SSR motif; (c) transferability of genomic and transcript SSR markers in four relatives of *B. rotunda*.

**Table 4.**
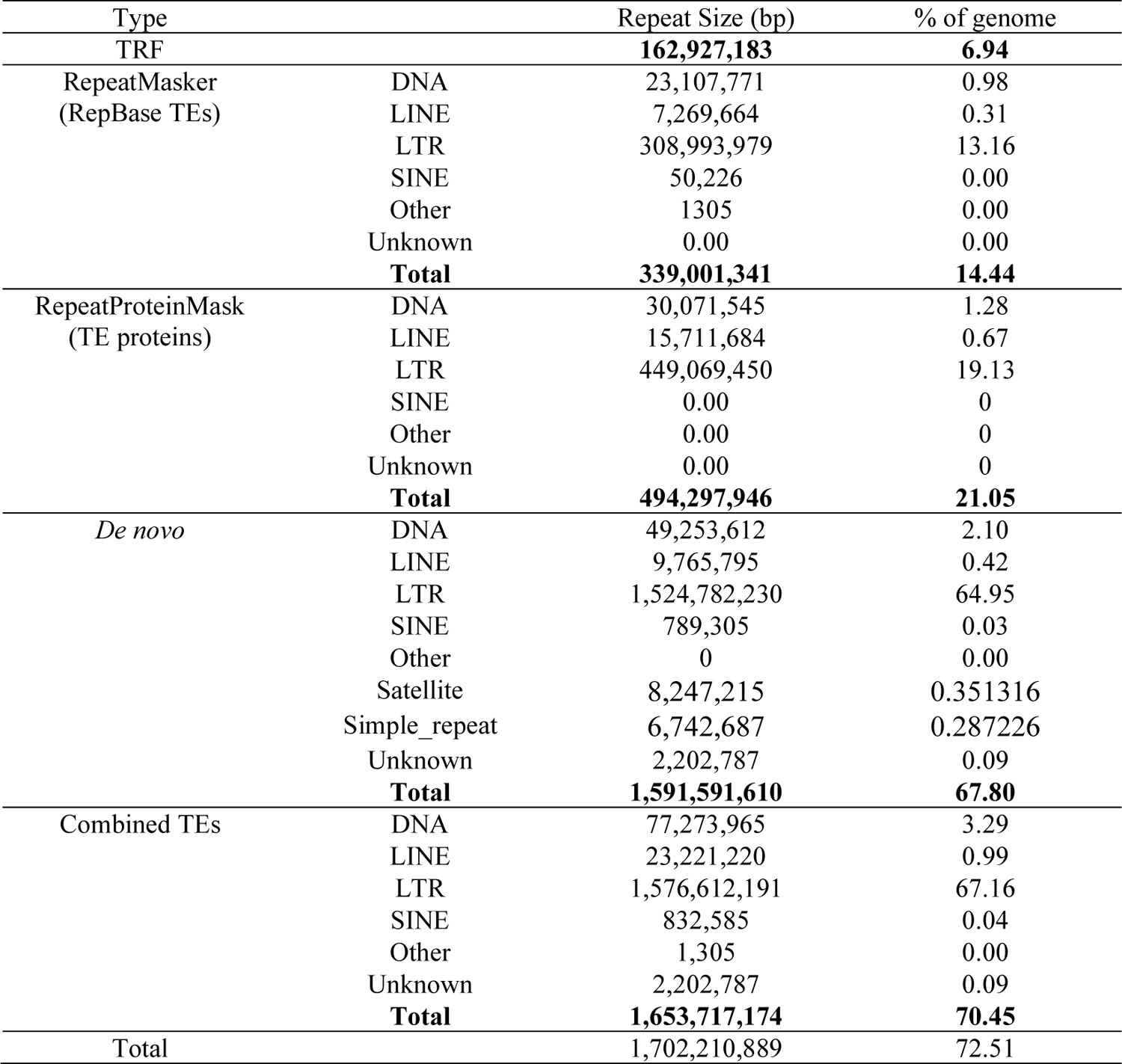
TEs Content in the assembled *B. rotunda* genome

**Table 5.** Genome and transcriptome-wide microsatellite identification and characterization in *B. rotunda*

| Item | Genome-wide | % | Transcriptome-wide | % |
| --- | --- | --- | --- | --- |
| Total number of sequences examined | 10627 |  | 95847 |  |
| Total size of examined sequences (bp) | 2347517452 |  | 66535881 |  |
| Total number of identified microsatellites | 238441 |  | 4579 |  |
| Number of microsatellites containing sequences | 10381 |  | 4032 |  |
| Sequences contain more than 1 microsatellites | 9803 |  | 384 |  |
| Microsatellites in compound formation | 4309 |  | 27 |  |
| Microsatellite's density (1 Microsatellites per ** bp) | 9845 |  | 14530 |  |
| Microsatellite's density (per Mbp) | 102 |  | 69 |  |
| Class I microsatellites | 82414 | 35.20 | 949 | 20.85 |
| Class II microsatellites | 151718 | 64.80 | 3603 | 79.15 |
| AT rich microsatellites | 176052 | 75.19 | 2778 | 61.03 |
| GC rich microsatellites | 43155 | 18.43 | 1275 | 28.01 |
| AT/GC balance microsatellites | 14925 | 6.37 | 499 | 10.96 |
| Mono-nucleotide repeats | 68961 | 28.92 | 1137 | 24.83 |
| Di-nucleotide repeats | 61439 | 25.77 | 574 | 12.54 |
| Tri-nucleotide repeats | 84932 | 35.62 | 2366 | 51.67 |
| Tetra-nucleotide repeats | 5330 | 2.24 | 148 | 3.23 |
| Penta-nucleotide repeats | 9917 | 4.16 | 185 | 4.04 |
| Hexa-nucleotide repeats | 7862 | 3.30 | 169 | 3.69 |
| Primer modelling was successful | 223678 | 93.81 | 3348 | 73.12 |
| Primer modelling failed | 14763 | 6.60 | 1231 | 36.77 |
| Non redundant primer | 132792 | 59.37 | 1888 | 56.39 |
| No of Primer Mapped on *Musa acuminata* genome | 100 | 0.075 | 30 | 1.59 |
| No of Primer Mapped on *Musa balbisiana* genome | 105 | 0.079 | 25 | 1.32 |
| No of Primer Mapped on *Musa Itinerans* genome | 102 | 0.077 | 32 | 1.69 |
| No of Primer Mapped on *Ensete ventricosum* genome | 121 | 0.091 | 27 | 1.43 |
| No of primer tested | 6 | 100 | 8 | 100 |
| No of primer amplified | 6 | 100 | 8 | 100 |

Mapping of *B. rotunda* SSR to close relatives using newly designed primer sequences showed that from the 93.81% of the genomic SSR and 73.12% of the transcriptome sequences suitable for SSR primer design, only a low number of primers mapped to the selected relatives, *Musa acuminata, Musa balbisiana, Musa itinerans* and *Ensete ventricosum* (Table 5). Overall, 224 G-SSR and 65 EST-SSR primers showed transferability into any of the four related species (with slightly more in Ensete), while only 42 genomic SSRs and 7 transcript SSRs were common to all five genomes (Fig. 2c, d). A subset of 14 *B. rotunda* SSR primer pairs (Table S3) were tested for their marker potentiality and showed that all amplified bands of the expected sizes for each species (Fig. S7).

The annotation of predicted protein-coding genes was a combination of homology-based and *de novo* prediction in addition to comparison with *B. rotunda* transcriptome data (Table S4). After consolidation, 73,102 protein-coding genes were predicted in the *B. rotunda* genome with an average transcript length of 4,312 bp (excluding UTR), CDS length of 1,360bp, average exon and intron lengths of 303bp and 812bp, and 4.49 exons per gene (Table S4). For the homology-based protein-coding gene predictions, protein sequences from four species (*M. acuminata*, *Phoenix dactylifera*, *Oryza sativa* and *Arabidopsis thaliana*) were mapped onto the *B. rotunda* genome. From these alignments, *B. rotunda* had the highest number of matches with *P. dactylifera* followed by *O. sativa*, *A. thaliana* and *M. acuminata* (Fig. S8). Functional annotation of the 73,102 predicted proteins from *B. rotunda* against seven databases enabled functional predictions for 97.8% of the predicted genes (Table 6). Non-coding RNA analysis of the assembly identified 213 microRNA (miRNA), 2,727 transfer RNA (tRNA), 486 ribosomal RNA (rRNA), and 2,136 small nuclear RNA (snRNA) genes (Table 7). A final genome annotation was performed by using MAKER together with *de novo* assembled non-redundant transcripts, predicted proteins, non-coding RNAs and repeats.

**Table 6.** Statistics of function annotation

| Database | Number | Percent (%) |
| --- | --- | --- |
| NR | 71,072 | 97.22 |
| InterPro | 69,525 | 95.11 |
| GO | 45,256 | 61.91 |
| KEGG | 59,649 | 81.60 |
| Swissprot | 57,622 | 78.82 |
| COG | 24,851 | 33.99 |
| TrEMBL | 70,990 | 97.11 |
| Total annotated | 73,102 | 97.81 |
| Unannotated | 1,602 | 2.19 |

**Table 7.** Non-coding RNA genes in the genome of *B. rotunda*

| Type | Copy | Average length (bp) | Total length (bp) | % of genome |
| --- | --- | --- | --- | --- |
| miRNA | 213 | 119 | 25,384 | 0.001081 |
| tRNA | 2,727 | 75 | 205,538 | 0.008756 |
| rRNA | 486 | 232 | 112,876 | 0.004808 |
| 18S | 105 | 666 | 69,922 | 0.002979 |
| 28S | 147 | 119 | 17,441 | 0.000743 |
| 5.8S | 40 | 148 | 5,931 | 0.000253 |
| 5S | 194 | 101 | 19,582 | 0.000834 |
| snRNA | 2,136 | 154 | 329,909 | 0.014054 |
| CD-box | 600 | 105 | 62,771 | 0.002674 |
| HACA-box | 53 | 134 | 7,091 | 0.000302 |
| splicing | 1,483 | 175 | 260,047 | 0.011078 |

### Functional classification by Gene Ontology

From a total of 95,847 unigenes derived from the *B. rotunda* transcriptome, 41,550 unigenes (43.35%) were found significantly scoring BLASTX hits against the NR protein database. Of these 6,850 (7.15% of the total unigenes) returned significant sequence alignments but could not be linked to any Gene Ontology entries; 6,038 (6.3%) of the GO mapped dataset did not obtain an annotation assignment and we could assign functional labels to 28,662 (29.9%) of the input sequences (Fig. S9). Species distribution among the BLASTX matches showed *M. acuminata* subsp. *malaccensis* to have a very high similarity score with 87,000 top BLASTX hits from *B. rotunda*. Other species matches included Ethiopian banana, *Ensete ventricosum* (Musaceae) with 70,000 hits, African oil palm, *Elaeis guineensis* (<u>Arecaceae</u>) with 62,500 BLASTX hits and date palm, *Phoenix dactylifera* (<u>Arecaceae</u>) with 62,000 BLASTX hits (Fig. S10). The annotated sequences assigned to GO classes based on Nr annotation in three clusters of biological process, molecular function and cellular component were categorized into 60 functional groups, with biological processes representing the largest number of sequences (Fig. 3a).

**Figure 3.**
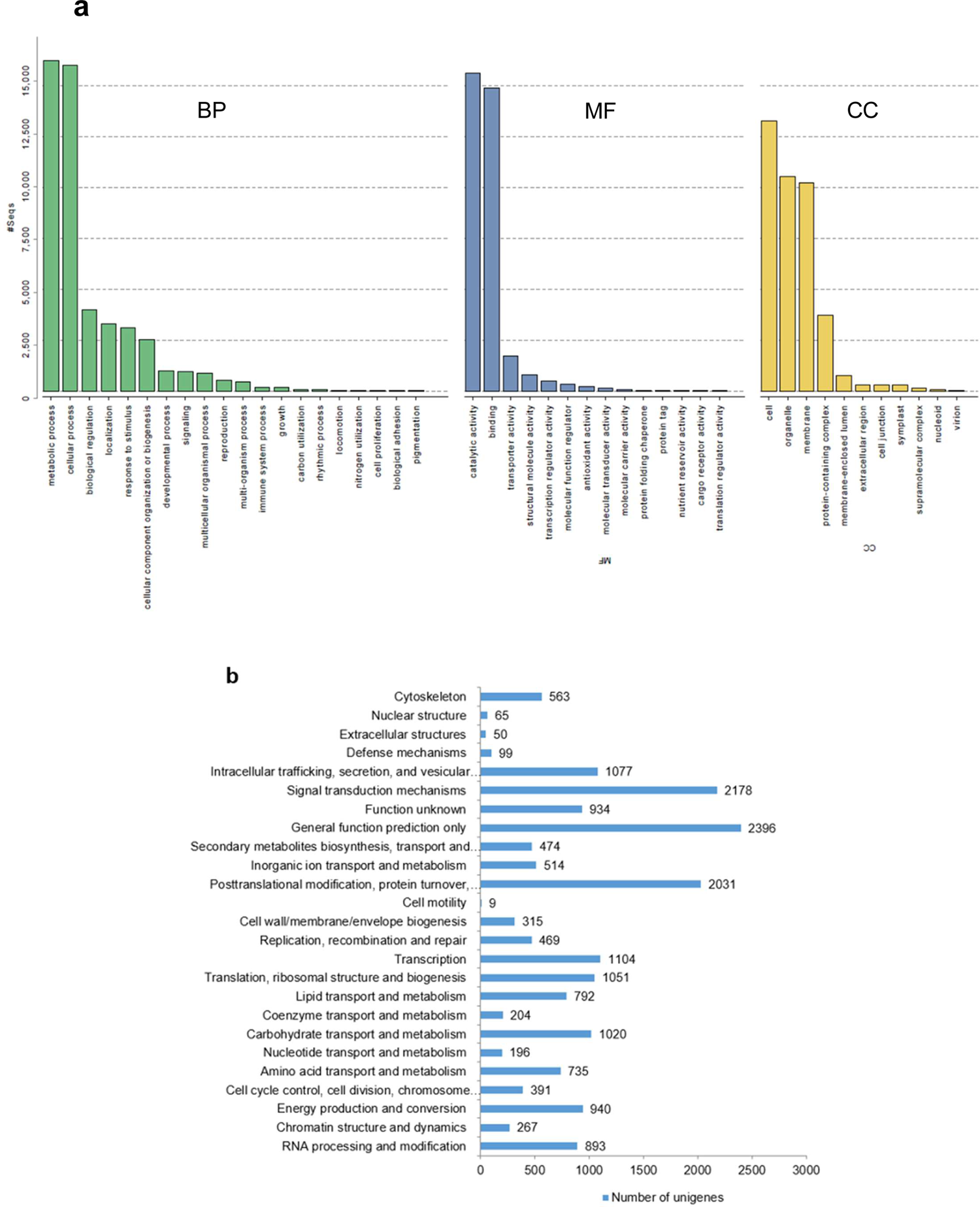
(a), Gene ontology (GO) classification of assembled unigenes of *B. rotunda*. Results are summarized in three main categories: biological process (BP), molecular function (MF), and cellular component (CC). The x-axis indicates the subgroups in GO annotation while the y-axis indicates the percentage of specific categories of genes in each main category; (b), Distribution of Eukaryotic Orthologous Groups (KOG) classification. A total of 18,767 assembled unigenes were annotated and assigned to 25 functional categories. The vertical axis indicates subgroups in the KOG classification and the x-axis represents the number of genes in each main category.

Blast2GO enzyme code (EC) annotation showed the distribution of *B. rotunda* predicted proteins among six main enzyme classes of oxidoreductases (1,400), transferases (3,500), hydrolases (2,250), lyases (450), isomerases (250), and ligases (270) (Fig. S11). The KOG function classification produced Nr hits for 18,767 unigenes which were annotated and classified functionally into 25 KOG functional categories including biochemistry metabolism, cellular structure, signal transduction, and molecular processing (Fig. 3b). The cluster for general function prediction represented the largest group with 2,396 genes followed by signal transduction mechanism (2,178) and posttranslational modification, protein turnover, and chaperons’ with 2,031 genes. All unigenes were analysed by comparison with the KEGG pathway database for further analysis of the *B. rotunda* transcriptome. Out of 28,662 annotated sequences, 1,494 (5.21%) unigenes were assigned to 145 predicted metabolic pathways.

### Phylogenetic orthology inference of *B. rotunda* genes

A total of 62,520 orthogroups were found with Orthofinder ^61^(Table S5) with matches of genes from *B. rotunda* to 979,315 genes from 13 other species (*Glycine max, Cucumis melo, Gossypium raimondii, Brassica napus, Arabidopsis thaliana, Solanum tuberosum, Solanum lycopersicum, Musa acuminata, Zea mays, Oryza sativa* subsp. *japonica, Hordeum vulgare, Phoenix dactylifera* and *Brachypodium distachyon*). Of these, 7,276 orthogroups were shared among all species and there were no single copy orthogroups (Table S5). The species tree inferred by STAG ^62^ and rooted by STRIDE ^63^ indicated that *B. rotunda* has the closest relationship with *M. acuminata* (order Zingiberales) and *P. dactylifera* (order Arecales) followed by members of the Poaceae family (*Z. mays*, *O. sativa* subsp. *japonica*, *H. vulgare*, and *B. distachyon*) and was distant from plant species from the Solanaceae, Brassicaceae, Malvaceae, Cucurbitaceae, and Fabaceae (Fig. 4a Table S5). UpSet plotting showed 7,276 orthogroups shared between *B. rotunda* and 13 selected reference genomes (Fig. 4b). 1,849 protein orthologs are specific for *B. rotunda* and 274 orthogroups shared among 13 selected reference genomes except for *B. rotunda*.

**Figure 4.**
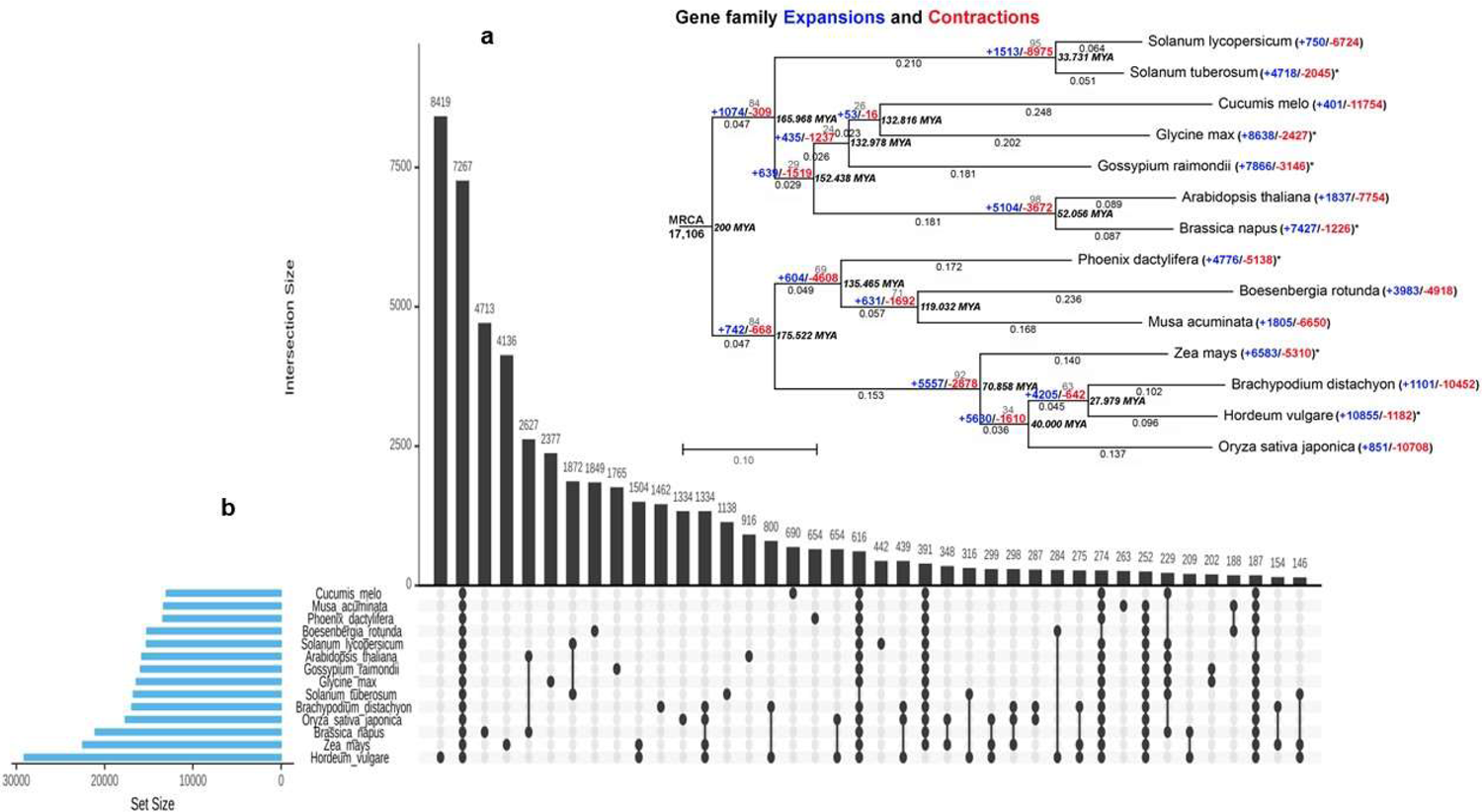
(a) Cross-genera phylogenetic analysis of *B. rotunda* and 13 other species; (b) UpSet plot showing unique and shared protein ortholog clusters of *B. rotunda* and 13 selected reference genomes. Connected dots represent the intersections of overlapping orthologs with the vertical black bars above showing the number of orthogroups in each intersection.

### Gene family expansion and contraction

Using the data generated from OrthoFinder ^61^, we explored gene family expansion and contractions in *B. rotunda* (Fig. 4a). In total, there are 17,106 gene families shared by the most recent common ancestor (MRCA). There were large numbers of gene families expanding (53– 10,855) or contracting (16–11,754) between 14 plant genomes (Fig. 4a). Our results show the substantial expansion of gene families in the Poaceae (5,557) followed by Brassicaceae (5,104) and the Pooideae subfamily (4,205). A large gene family contraction was observed in Solanaceae (8,975). Interestingly, the majority of the genomes with reported ancient whole genome duplication or massive segmental duplications or major chromosomal duplications show higher number of gene family duplications than gene family losses (indicated by asterisks in Fig. 4a).

### Transcriptome changes of *B. rotunda* unigenes related to flavonoid and phenylpropanoid biosynthesis pathways

Transcriptome analysis showed in total 167 unigenes from *B. rotunda* were mapped to five different classes of enzymes including oxidoreductase, transferase, ligase, lyase, and hydrolase in flavonoid and phenylpropanoid pathways. Of these, only 23 enzymes showed differential expression in the different samples i.e., *in vitro* leaf (IVL), embryogenic callus (EC), and non-embryogenic calli (dry callus (DC) and watery callus (WC)) using *ex vitro* leaf (EVL) samples as the comparator (Fig. 5, Table S6). The first enzyme in the phenylpropanoid pathway is phenylalanine ammonia-lyase (PAL) which converts phenylalanine to cinnamic acid. PAL was expressed at the lowest levels among all samples in IVL with the highest expression level in WC (indicated by dark red squares in Fig. 5). Then coenzyme A (CoA) will be attached to cinnamic acid or *p*-coumaric acid by 4-coumarate–CoA ligase (4CL) and form cinnamoyl-CoA or *p*-coumaroyl-CoA. This enzyme showed relatively higher expression in all samples except EC. In the phenylpropanoid pathway, cinnamic acid is also converted to coumarinate by Beta-glucosidase (BGLU) to produce coumarin. BGLU was expressed in all samples, with the highest expression level in non-embryogenic calli (NEC). Then CHS, chalcone synthase (CHS) converts cinnamoyl-CoA to pinocembrine chalcone and p-coumaroyl CoA to naringenin chalcone. CHS was expressed in all samples except IVL with the highest expression level in WC. In the next step, the two flavanones of pinocembrin and naringenin are synthesised by chalcone isomerase (CHI). CHI was expressed in all samples except IVL with the highest expression level in EC. Pinocembin is converted to pinostrobin by flavanone-3-hydroxylase (F3H) which serves as precursor of panduratin A synthesis. Expression analysis of unigenes related to F3H enzyme and dihydroflavonol 4-reductase (DFR) which are involved in the synthesis of anthocyanidins such as pelargonidin, cyanidin, and delphinidin, showed DFR to be more highly expressed in DC and WC compared to other samples, while F3H was only relatively up-regulated in WC. Other enzymes in the phenylpropanoid pathway include hydroxycinnamoyl-CoA shikimate (HCT), cinnamoyl-CoA reductase (CCR), cinnamyl alcohol dehydrogenase (CAD; EC1.1.1.195), caffeoyl-CoA O-methyltransferase (CCOAOMT), and lactoperoxidase enzyme (LPO) involved in monolignols synthesis such as *p*-hydroxyphenyl (H), guaiacyl (G) and syringyl (S). Among them, HCT, CCOAOMT, and CAD showed higher expression in all samples, except IVL for CAD, while CCR showed higher expression in WC. The gene expression differences between the tissue samples for cinnamic acid 4-hydroxylase (C4H), ρ-coumarate 3-hydroxylase (C3H), ferulate 5-hydroxylase (F5H), caffeic acid O-methyltransferase (COMT), and cinnamyl alcohol dehydrogenase (CAD) were below the threshold of FPKM without any differential expression in studied samples.

**Figure 5.**
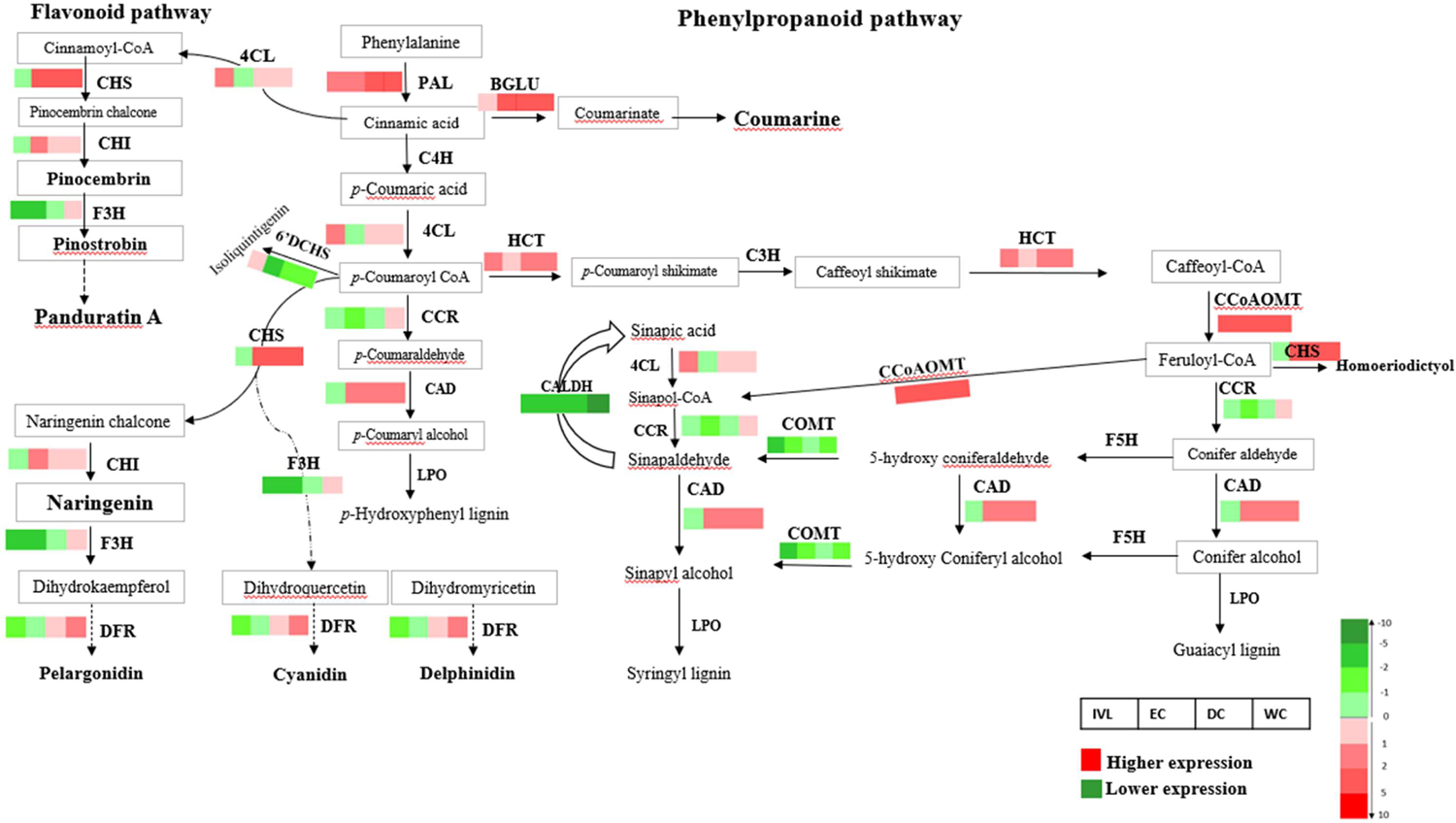
Scheme of the flavonoid and phenylpropanoid biosynthetic pathways in *B. rotunda* based on KEGG pathways. Genes encoding enzymes for each step are indicated as follows: CAD, cinnamyl alcohol dehydrogenase; and BGLU, Beta-glucosidase; CALDH, coniferyl-aldehyde dehydrogenase; C4H, cinnamic acid 4-hydroxylase; 4CL, 4-coumarate–CoA ligase; CHS, chalcone synthase; CHI, chalcone isomerase; CCoAOMT, caffeoyl-CoA 3-O-methyltransferase; C3H, ρ-coumarate 3-hydroxylase; CCR, cinnamoyl-CoA reductase; COMT, caffeic acid O-methyltransferase; 6’DCHS, 6′-deoxychalcone synthase; DFR, dihydroflavonol 4-reductase; F3H, flavonoid 3-hydroxylase; F5H, ferulate 5-hydroxylase, HCT, Hydroxycinnamoyl-CoA shikimate; LPO, Lactoperoxidase; PAL, phenylalanine ammonia lyase. Beside each enzyme, four boxes shown (from left to right): *In vitro* leaf (IVL), Embryogenic callus (EC), Dry callus (DC), Watery callus (WC). Red boxes indicate relatively higher mRNA expression compared to the leaf sample with the highest levels in darker red. Green boxes indicate relatively lower expression compared to the leaf sample. The colour box is based on log_2_FC values.

### DNA methylation analysis using bisulfite sequencing

Genome wide methylation percentages determined from bisulfite sequence data from leaf and four tissue cultured samples, were higher in all methylated cytosine contexts for samples from EVL (CpG 73.2%, CHG 36.2%, CHH 33.7%) and IVL (CpG71.3%, CHG 35.4%, CHH 33.5%). The lowest levels were for EC (CpG 53.4%, CHG 18.5%, CHH 25.3%), followed by WC (CpG 63.8%, CHG 21.9%, CHH 28.1%) and DC (CpG 68.4%, CHG 25.9%, CHH 28.6%). We also evaluated DNA methylation levels of three groups of genes (30 genes in total) including DNA methyltransferase-related genes across the genome of *B. rotunda* (Fig. 6a-d). In general, CHH methylation levels were higher than methylation levels in CpG and CHG context and out of 30 genes, 22 genes (73.3%) showed low methylation levels (<0.1) in CHG and CHH cytosine contexts whereas only 30% of the genes showed low methylation levels in the CpG context. Cytosine methylation of methylation-related genes in all cytosine contexts (CpG, CHG & CHH) was the highest for *DRM2* and followed by *MET1*, and *CMT3* (Fig. 6a-d). Among somatic embryogenesis-related genes, *WOX* gene was heavily methylated in CpG and CHG contexts compared to other somatic embryogenesis-related genes (*LEC2, BBM, SERK*) (Fig. 6a-d). For pathway-related genes, *LPOs* methylated more in all studied samples compared to other genes.

**Figure 6.**
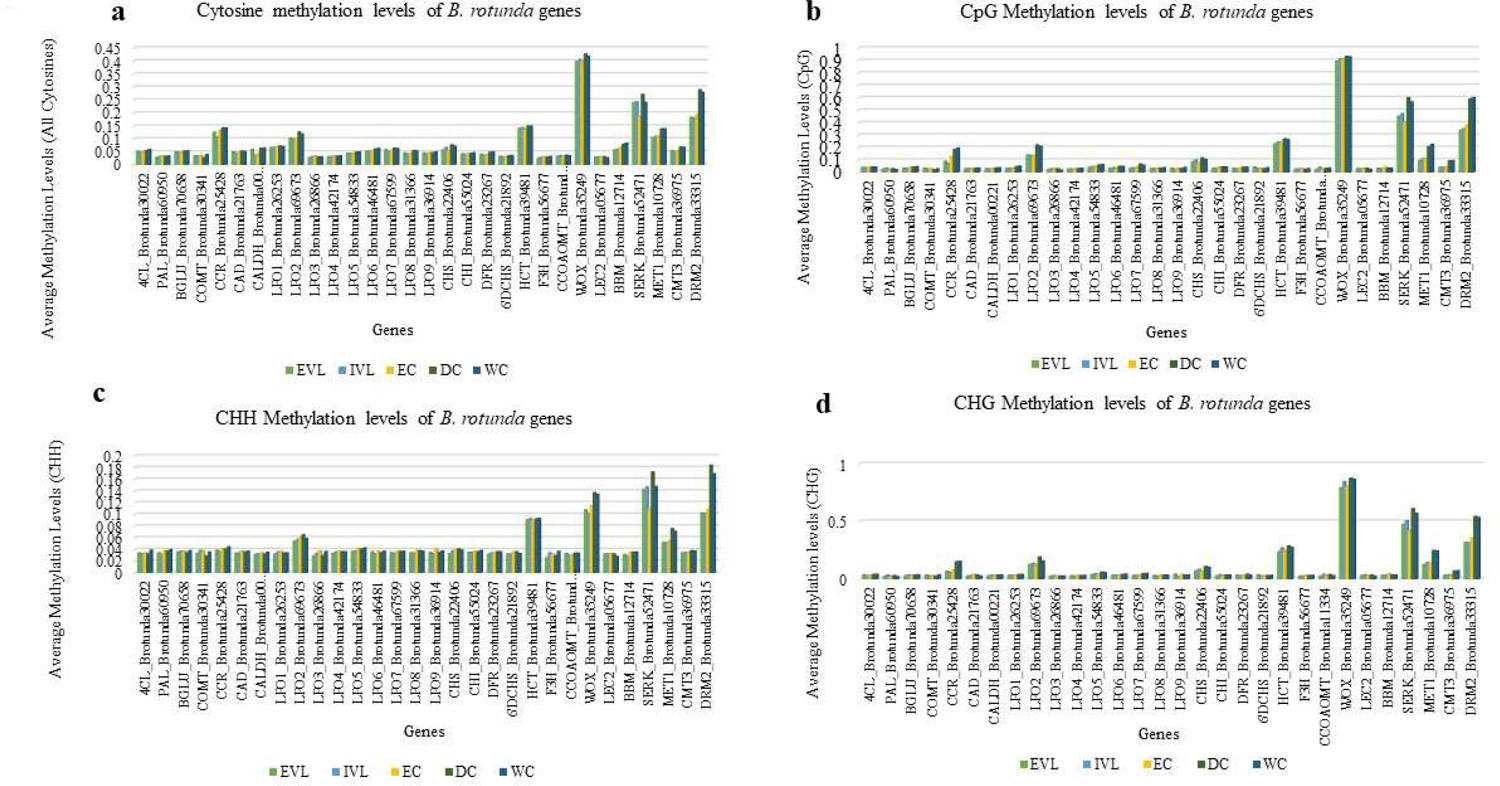
(a-d) Average methylation levels of DNA methyltransferase-related genes (MET1, CMT3, DRM2), somatic embryogenesis genes (SERK, BBM, LEC2, and WUS), and genes involved in flavonoid and phenylpropanoid biosynthesis pathways in different samples of *B. rotunda*; *ex-vitro* leaf (EVL), *in vitro* leaf (IVL), Embryogenic callus (EC), Dry callus (DC), Watery callus (WC). a) cytosine methylation; b) CpG methylation; c) CHG methylation; and d) for CHH methylation. CAD, cinnamyl alcohol dehydrogenase; and BGLU, Beta-glucosidase; CALDH, coniferyl-aldehyde dehydrogenase; C4H, cinnamic acid 4-hydroxylase; 4CL, 4-coumarate–CoA ligase; CHS, chalcone synthase; CHI, chalcone isomerase; CCoAOMT, caffeoyl-CoA 3-O-methyltransferase; C3H, ρ-coumarate 3-hydroxylase; CCR, cinnamoyl-CoA reductase; COMT, caffeic acid O-methyltransferase; 6’DCHS, 6′-deoxychalcone synthase; DFR, dihydroflavonol 4-reductase; F3H, flavonoid 3-hydroxylase; F5H, ferulate 5-hydroxylase, HCT, Hydroxycinnamoyl-CoA shikimate; LPO, Lactoperoxidase; PAL, phenylalanine ammonia lyase; WOX, Wuschel; LEC3, Leafy cotyledon 2; BBM, Baby boom; SERK, Somatic embryogenesis receptor-like kinase; MET1, Methyltransferase 1; CMT3, Chromomethylase 3; DRM2, Domain rearranged methyltransferase 2.

### Correlation between gene expression levels and DNA methylation levels of genes related to methylation, somatic embryogenesis and secondary metabolite pathway

From gene expression analysis, we observed that the expression level of DNA methyltransferase genes, *MET 1, CMT 3, and DRM2* was higher in callus than leaf samples and was highest in embryogenic callus (EC) for all three genes and lowest in *in vitro* leaf (IVL) (Fig. 7a). *DRM2* showed the lowest level of expression and the highest level of DNA methylation. DNA methylation levels of these genes at CpG, CHG, CHH cytosine contexts were the highest in DC and WC with similar and lower methylation levels in the embryogenic callus and leaf samples. Overall, expression of methylation-related genes was higher in samples EC, DC, and WC but other than for *CMT3*, which showed an inverse relationship between expression level and methylation levels, there was no clear correlation between level of DNA methylation and level of gene expression (Fig. 7b). Similarly, while there were different expression patterns for the four somatic embryogenesis-related genes *SERK*, *BBM*, *LEC2*, and *WOX* between different leaf and callus samples (Fig. 7c), the DNA methylation level of each gene across the different leaf and callus samples was largely unchanged (Fig. 7d). A comparison of 23 of *B. rotunda* genes involved in flavonoid and phenylpropanoid pathways showed them to be expressed differentially in *B. rotunda* leaf and callus samples. Among them, *BGLU*, *CAD*, *CHS*, *LPO8*, *LPO9* and *PAL* were expressed more highly in callus than in leaf samples (Fig. 7e). The highest level of DNA methylation was observed for *HCT, CCR*, and *LPO2* genes in all studied samples and again, there was no general correlation between gene expression levels and methylation levels for these samples (Fig. 7f).

**Figure 7.**
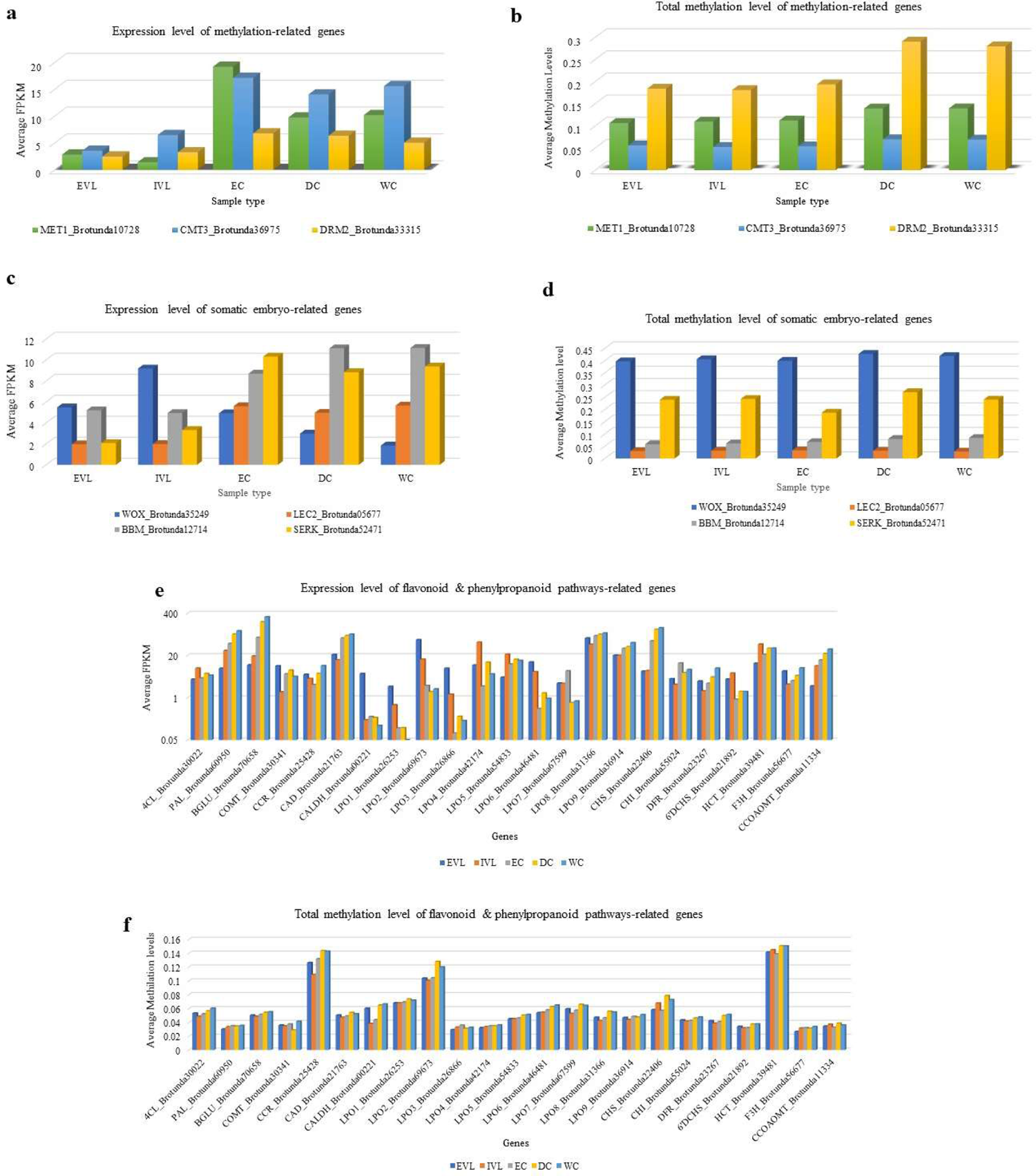
Expression level and total methylation level in all cytosine contexts of methylation-related genes (a & b), somatic embryogenesis-related gene (c & d) and flavonoid and phenylpropanoid biosynthesis pathways-related genes (e & f) in *ex vitro* leaf (EVL), *in vitro* leaf (IVL), embryogenic callus (EC), dry callus (DC), and watery callus (WC). CAD, cinnamyl alcohol dehydrogenase; and BGLU, Beta-glucosidase; CALDH, coniferyl-aldehyde dehydrogenase; C4H, cinnamic acid 4-hydroxylase; 4CL, 4-coumarate–CoA ligase; CHS, chalcone synthase; CHI, chalcone isomerase; CCoAOMT, caffeoyl-CoA 3-O-methyltransferase; C3H, ρ-coumarate 3-hydroxylase; CCR, cinnamoyl-CoA reductase; COMT, caffeic acid O-methyltransferase; 6’DCHS, 6′-deoxychalcone synthase; DFR, dihydroflavonol 4-reductase; F3H, flavonoid 3-hydroxylase; F5H, ferulate 5-hydroxylase, HCT, Hydroxycinnamoyl-CoA shikimate; LPO, Lactoperoxidase; PAL, phenylalanine ammonia lyase.

## Discussion

We present a genome assembly of *Boesenbergia rotunda* (2n=36) with an estimated genome size of 2.4Gb. The genome of the plant we sequenced, when in cultivation a largely vegetatively propagated species, shows an unusually high heterozygosity of 3.01%, suggesting that the cultivar may be of hybrid origin or may have undergone whole genome duplication events. This is also suggested based on the large number of unigenes in *B. rotunda*, notably more than twice that of *Ensete glaucum* ^56^, and 46,765 duplication events (65.8% of the *B. rotunda* genome, with at least 50% support). As noted in *Citrus limon* ^64^, high levels of heterozygosity complicate the assembly process. Due to the clonal propagation nature of the fingerroot ginger, offspring resulting from the sexual hybridization is rather limited. Thus, we applied a similar approach as reported by Chin et al. (2016) and Baek et al. (2018), for the assembly of the *B. rotunda* genome ^65, 66^. The sequencing assembly of *B. rotunda* using long PacBio reads, in addition to the Illumina short-reads, and followed by assembly using FALCON assembler resulted in a scaffold number of 10,627. The relatively high scaffold number is not unexpected considering the high repeat content (72.51%) of the *B. rotunda* genome, coupled with the relatively high level of heterozygosity (3.01%), and the lack of any molecular marker and breeding data for *B. rotunda*. Future mapping and marker studies could help to resolve an assembly into the anticipated 18 chromosomes, as could more recent technologies such as single chromosome sequencing and optical mapping ^55^.

Sequence information for other *Boesenbergia* species is not yet available, with the closest relative of *B. rotunda* from sequenced genomes at the time of our study being *M. acuminata*, based on previous analyses using amino acid data from single genes including chalcone isomerase (CHI) ^67^ and phytyltransferase (BrPT2) ^68^. Our phylogeny analysis also showed *M. acuminata* as the closest relative among those compared, with *Z. mays*, *O. sativa*, *H. vulgare*, and *B. distachyon* from the Poaceae family, more distantly related, as expected.

The repeat content of the *B. rotunda* at ∼72% of the assembled genome is high compared to many other plant genomes in this order such as *Musa itinerans* (38.95%) ^69^ and *M. acuminata* (35.43%) ^70^, but similar to that of *Z. officinale* (ginger official) at 81% ^71^. A higher level of repeat content has been observed to correlate with larger genome sizes in the Fabaceae ^72^ and *Melampodium* ^73^. Both of those reports suggest the greater genome size to be largely driven by Ty3/gypsy LTR-retrotransposons and it is interesting to note that *B. rotunda* also has a high LTR content of 64%. While data for genome sizes and content are not yet available for other Boesenbergia species, the *Z. officinale* genome has a similar high value of 61% LTR which was also suggested to contribute to the high genome size ^74^. Studies in other plant species reported that plant genomes generally have over 50% transposable elements content (e.g., maize) while some small plant genomes such as Arabidopsis may have as low as 10% repeat content ^75–77^. Cytosine methylation is usually much denser in transposons than in genes ^78–80^ and this has also been correlated with evolution of genome size in angiosperms ^76^. The large genome size and high repeat content of *B. rotunda* with relatively low gene body cytosine methylation levels of the genes selected for observation in the current study, fit well with this model and it will be interesting to compare this with other Boesenbergia species in the future when similar data becomes available.

As DNA methylation is dynamic, we saw variations in global DNA methylation levels in the different samples. Unmethylated DNA has been shown to demarcate expressed genes ^81^ and so to be able to examine this in the context of gene expression in *B. rotunda* and to add depth to our genome data, we included deep sequencing of leaf and callus transcriptomes from *B. rotunda*. There are several alternative tools for the *de novo* assembly of RNA-seq short reads into a reference transcriptome and we compared analysis from four assemblers. The quality of assembly was noticeably affected by both k-mer size and assembler tool, with Oases delivering the highest N50 size and average contig length at k-mer 21 compared to at k-mer 24 or other assemblers (Figure S4, Table 3), indicating more effective and accurate assembly. In comparison to a previous transcriptome assembly of *B. rotunda* by SOAPdenovo-Trans *de novo* assembler, our study obtained a longer N50 size (1,019) compared to an N50 value of 236 reported by ^40^. An Oases assembly of genome sequence data from a Fern, *Lygodium japonicum* was also found to give the best mean transcript length and N50 size when compared to assemblies using Trinity and SOAPdenovo-Trans ^82^. The BUSCO assessment of *B. rotunda* transcriptome data also showed that Oases had higher numbers of complete and single copy contigs and less fragmented contigs. Based on this, the transcriptome assembly using Oases offered an improved resource for genome annotation and the gene expression study in *B. rotunda*.

We focused functional aspects of the *B. rotunda* genome study on the methylation and the flavonoid and phenylpropanoid pathways, as the chalcone, panduratin A, is considered one of the most promising bioactive compounds from *B. rotunda* and previous studies from our research group had indicated DNA methylation may influence gene expression in tissue cultured materials ^83, 84^. From the 23 flavonoid and phenylpropanoid pathway genes that showed differential expression between leaf and any of the callus samples, most were more highly expressed in EC, DC, and WC, including *PAL*, *CHS*, *CHI*, *DFR*, *BGLU*, *HCT*, *CCOAOMT*, and *CAD* (Fig. 7) with highest expression level in the non-embryogenic callus (DC and WC). This aligns with previous Ultra Performance Liquid Chromatography-Mass Spectrometry (UPLC-MS) data showing WC followed by DC to have a higher concentration of panduratin, pinocembrin, pinostrobin, cardamonin and alpinetin ^42^. Based on this, the unigenes identified in the genome assembly that correspond to CHS and CHI, encode key enzymes in the biosynthesis of panduratin A in *B. rotunda*. Although DNA methylation plays an important role in the regulation of gene expression, comparison of the methylation of the differentially expressed flavonoid and phenylpropanoid pathway genes, with their cytosine methylation showed no obvious patterns to indicate any correlation for this gene set.

As our samples included embryogenic and non-embryogenic callus tissue, we also evaluated the expression level of DNA methylase genes (*MET1*, *CMT3*, *DRM2*) and genes related to somatic embryogenesis (*SERK*, *BBM*, *LEC2*, *WUS*) with DNA methylation levels across the genome of *B. rotunda* based on bisulfite sequence analysis. An earlier study with some quantitative qRT-PCR validation suggested that the higher level of expression of methyltransferase-related genes and the lower CG, CHG and CHH sequence contexts in EC samples was negatively correlated with the total methylation level of DNA methyltransferase-related genes ^84^. We did observe a similar pattern for *CMT3* in all five sample types in the current study (Fig. 7), however, no similar correlation between expression level and cytosine methylation was observed in the current data for the other genes examined. The lack of correlation between transcript expression and the respective gene body methylation from our data may be due to the limitations of the current genome assembly such that the cis regions could not be well annotated. In the future a higher resolution genome assembly for *B. rotunda* would be useful to examine the methylation data from the current study.

Although only a minor portion of the *B. rotunda* genome at around 0.35%, microsatellites are key elements in plant genomes. Among these, short sequence repeat microsatellites (SSRs) have found wide utility as co-dominant markers useful in breeding and diversity studies ^85, 86^. In this study, we identified genomic and EST-SSRs from *B. rotunda*, designing primers and showing several to have transferability to Musa and Ensete genomes, mostly *in silico* analysis, but with 14 tested in PCR experiments. Boesenbergia, Musa and Ensete are members of the same plant family *Zingiberales*, and all have abundant AT-rich SSR sequences, however they are not from the same genus, so are phylogenetically somewhat distanced as reflected in the fairly low numbers with potential as markers across these species. Nevertheless, these newly developed SSR markers enhance the genetic resources for *B. rotunda* as well as the plant family *Zingiberales* and these markers could be utilized for genotyping, population structure analysis, association studies, cultivar identification as well as any other breeding application of the *Boesenbergia spp*.

In conclusion, the genome assembly of *B. rotunda* covers some 2,300 Mbp divided among 18 relatively similar submetacentric chromosomes. The cultivated accession sequenced was highly heterozygous. The genome assembly, transcriptome, gene expression, SSR analysis and DNA methylation data from this study are resources that will allow further understanding of the unique secondary metabolite properties and their biosynthetic pathways in the genus Boesenbergia and for functional genomics of *B. rotunda* characteristics, evolution of the ginger plant family and potential genetic selection or improvement of gingers.

## Materials and methods

### Ethics

The conduct of this research was approved by the grant management committee of the University of Malaya, headed by the Director of the Institute of Research Management and Monitoring, Professor Noorsaadah Abdul Rahman and did not involve the use of any human, animal, or endangered or protected plant species as materials.

### Plant materials and establishment of *in vitro* samples

Rhizomes of *B. rotunda* (L.) Mansf. were obtained from a commercial farm in Temerloh, Pahang, Malaysia (Latitude: 3.27° N, Longitude: 102.25° E) and propagated in the laboratory to generate all sample materials following methods described by Karim et al. (2018b) ^84^. Initially, the plants were washed thoroughly under running tap water for 10 min, then air dried for 30 min before insertion into black polybags to promote sprouting. Samples were sprayed with water every day to induce growth of shoots and leaves. The samples included young *ex vitro* leaf (EVL) samples, collected from rhizome-derived plants at four weeks after potting; Callus samples cultured from meristematic block explants subcultured on MS medium supplemented with 30 g L-1 sucrose and 2 g L-1 Gelrite® with 2,4-dichlorophenoxy acetic acid (2,4-D) at concentrations of 1 mg L-1 (4.5 µM) for watery callus (WC), 3 mg L-1 (13.5 µM) for embryogenic callus (EC) and 4 mg L-1 (18 µM) for dry callus (DC). The WC, EC and DC samples were collected after four weeks on the respective media (8 weeks after initial culturing from explant). *In vitro* leaves (IVL) from plants regenerated from embryogenic calli placed on regeneration media (MS0) were collected after 8 weeks (16 weeks after initial culturing from meristematic block explants) ^83^.

### DNA extraction and sequencing for genome and bisulfite sequence (BS-seq) analysis

Total genomic DNA was extracted using a modified cetyl trimethyl ammonium bromide (CTAB) method from *ex vitro* leaf (EVL) of *B. rotunda* ^87^. The quality and quantity of extracted DNA were determined by measuring the absorbance at A260nm and A280nm using a NanoDrop 2000 Spectrophotometer (Thermo Fisher Scientific Inc., Waltham, MA, USA) and Qubit® 2.0 Fluorometer (Thermo Fisher Scientific Inc., Waltham, MA, USA). The DNA sample was sent to BGI Shenzhen (Shenzhen, China) for library construction and *de novo* sequencing on the Illumina HiSeq2000 and HiSeq2500 platform (Illumina Inc., San Diego, CA, USA) and the PacBio RS II platform (PacBio Inc., CA, USA). Different insert size (bp) libraries were prepared using the best quality DNA samples with an A260nm/A280nm ratio between 1.7–1.9. For library construction, DNA was fragmented, end repaired, 3’A tailed, adapter ligated, and amplified by PCR ^88^. For BS sequence analysis, genomic DNA of *B. rotunda ex vitro* leaf (EVL), embryogenic callus (EC), dry callus (DC), watery callus (WC), and *in vitro* leaf of regenerated plants (IVL) were sequenced after being treated by sodium bisulfite. The sequencing was carried out by a commercial service provider, Sengenics Sdn. Bhd., Malaysia. A total of five samples (three biological replicates for each of five samples) were sequenced to generate paired-end reads using an Illumina HiSeqTM 2000 platform (Illumina Inc., San Diego, CA, USA) according to the manufacturer’s instructions.

### RNA extraction and sequencing for transcriptome (RNA-seq) analysis

Total RNA was isolated from *ex vitro* leaf (EVL), embryogenic callus (EC), dry callus (DC), watery callus (WC), and *in vitro* leaf of regenerated plants (IVL) using a modified cetyl trimethyl ammonium bromide (CTAB) method ^89^. Three independent rounds of RNA (n = 3) were prepared for each sample. Total RNA was measured using a NanoDrop 2000 Spectrophotometer (Thermo Fisher Scientific Inc., Waltham, MA, USA) and RNA integrity was determined using an Agilent 2100 Bioanalyzer (Agilent Technologies Inc., CA, USA). RNA samples with absorbance ratios A260nm/A280nm ranging from 1.8 to 2.2, and an A260nm/A230nm ratio higher than 1.0 and an RNA integrity number (RIN) higher than 7.0, were sent to BGI-Shenzhen (Shenzhen, China) for library construction and sequencing on the using Illumina Genome Analyzer IIx (GAIIx) platform (Illumina Inc., San Diego, CA, USA) to generate single-end reads.

### Determination of chromosome number and location of 45S and 5S rDNA sites on metaphase chromosomes of *B. rotunda (*2n=36) using fluorescent in situ hybridization (FISH)

The FISH procedure was adapted to Schwarzacher and Heslop-Harrison (2000)^90^. ‘Fingers’ of *B. rotunda* were placed in shallow dishes with soil to initiate root growth and kept in the glasshouse at the University of Leicester, UK. Newly grown roots tips of 1-2cm length were treated with 2 mM 8-hydroxyquinoline at growth temperature for 2 hours followed by incubation overnight at 4°C, and then fixed with 96% ethanol:glacial acetic acid (3:1). Roots were digested for 1-3h at 37°C with a mixture of cellulose (32U/ml, Sigma-Aldrich C1184), ‘Onozuka’ RS cellulose (20U/ml), pectinase (from Aspergillus niger, Sigma-Aldrich P4716) and Viscozyme (20U/ml, Sigma-Aldrich V2010) in 10mM citric acid/sodium citrate buffer (pH4.6). Chromosome preparations of dissected meristems were made in 60% acetic acid by squashing under a cover slip. Slides were stored at −20°C until FISH.

The 45S rDNA and 5S rDNA probe were labelled by random priming (Invitrogen) with digoxigenin 11-dUTP or biotin 11-dUTP (Roche) using the linearised clone pTa71 (from *Triticum aestivum*, Gerlach and Bedbrook 1979) or the PCR amplified insert of clone pTa794 (from *T. aestivum*, Gerlach and Bedbrook 1979), respectively ^91^. For hybridization, 50-100ng of labelled probe were prepared in 40-50µl mixture of 40% (v/v) formamide, 20% (w/v) dextran sulphate, 2x SSC (sodium chloride sodium citrate), 0.03μg of salmon sperm DNA, 0.12% SDS (sodium dodecyl sulphate) and 0.12mM EDTA (ethylenediamine-tetra acetic acid). Chromosomes and probe mixture were denatured together at 70°C for 6-8 mins, before cooling down slowly to 37°C and hybridized for 16h at 37°C. Slides were washed at 42°C in 0.1xSSC and hybridization sites were detected with anti-digoxigenin-FITC (2µg/ml; Roche) and Streptavidin-Alexa594 (1µg/ml; Molecular Probes). Chromosomes were counterstained with DAPI (4’,6-diamidino-2-phenylindole, 4µg/ml) and mounted in CitifluorAF. Slides were examined with a Nikon Eclipse 80i microscope and images were captured using NIS-Elements v2.34 (Nikon, Tokyo, Japan), and a DS-QiMc monochrome camera. Images were pseudocoloured and final figures were prepared with Adobe Photoshop CC2018 using enhancements that treat all pixels of the image ^90^.

### k-mer analysis for genome size estimation

The genome size of *B. rotunda* was estimated based on the flow cytometry and K-mer analysis. We determined the genome size (G) of *B. rotunda* as an unknown sample with flow cytometry on a MACSQuant Analyzer (Miltenyl Biotec Inc., BG, Germany), using soybean (*Glycine max* cv. *Polanka* (G) 2C = 2.50 pg DNA) and Pea (*Pisum sativum* cv. Ctirad (P) 2C = 9.09 pg DNA) as internal standards and propidium iodide as the stain. Each plant (sample and comparator) was compared using an average of four biological replicates ^52, 92, 93^. We also performed K-mer analysis to estimate the *B. rotunda* genome size and heterozygosity rate using Jellyfish ^94^ and GenomeScope ^95^.

### Genome assembly

A combination of sequencing technologies of PacBio RSII platform, Illumina HiSeq 2500 paired-end reads (PE) with 450bp insert size library, and Illumina HiSeq 2000 mate-pair reads (MP) with insert size libraries of 2, 5, 10, 20, and 40kb was performed for genome assembly. Before assembly, Illumina HiSeq sequence reads were filtered by removing adaptors and low-quality nucleotides. PacBio reads were filtered to remove the short reads of less than 500bp or a quality score lower than 0.8, then error correction for the long reads done by FALCON ^96^, following the general principles proposed by ^97^. We have tried to use several *de novo* assemblers to construct the assembly with both Illumina and PacBio reads. Finally, we chose the SMARTdenovo ^98^. Corrected PacBio reads were assembled with SMARTdenovo software (https://github.com/ruanjue/smartdenovo) to construct contigs. For PacBio data, constructed contigs were subsequently polished by stand-alone consensus modules, ^97^ and Pilon software ^99^ for Illumina PE reads. Polished contigs were used as input for scaffolding. Scaffolds were constructed by SOAP scaffolding, SSPACE tool ^100^ with Illumina mate-pair reads (2k-40k) with default parameters to extend the length of scaffolds for the raw assembly. The gaps within scaffolds, consensus sequences generated from PacBio sub-reads were filled using PBJelly2 ^101^. Finally, the scaffolds were corrected by Pilon ^99^ with Illumina PE reads to correct the assembly errors and obtained final genome assembly.

### Genome assembly quality assessment

The completeness of the assembly was tested by searching for 1440 core eukaryotic genes using Benchmarking Universal Single-Copy Orthologs (BUSCO) (v2.0) ^60^. To assess the quality of the genome assembly, the Illumina paired-end 250bp read data was mapped to the contig using BWA-MEM (version 0.7.15-r1142) ^102^.

### Repeat annotation

Tandem repeats were identified with tandem repeat finder (TRF) ^103^ (version 4.0.4). Transposable elements (TE) were identified with integrated homology-based and *de novo* methods ^52^. Homology-based prediction was done at the DNA and protein levels by comparing the assembly to the RepBase v.20.04 ^104^ database as a query library using RepeatMasker v.4.0.7 (http://www.repeatmasker.org/) and ProteinRepeatMask v.4.0.7 (http://www.repeatmasker.org/). To search those absent TEs in RepBase library, *de novo* repeat library was constructed using RepeatModeler v.1.0.10 (http://www.repeatmasker.org/) to run against *B. rotunda* genome assembly using RepeatMasker v.4.0.7 (http://www.repeatmasker.org/).

### Gene annotation

Three approaches were employed in gene prediction: Homolog, *de novo*, and RNA-Seq. For generation of homology-based predictions, the gene sets from four species i.e. *M. acuminata* (http://www.promusa.org/Musa+acuminata), *P. dactylifera*, *O. sativa* (http://rice.plantbiology.msu.edu/) and *A. thaliana* (https://www.arabidopsis.org/) were downloaded. The nonredundant protein sequences for each gene set was searched by TBLASTN. For generation of expression-based evidence, RNA-seq short reads originating from *ex vitro* leaf (EVL), *in vitro* leaf (IVL), embryogenic callus (EC), dry callus (DC) and watery callus (WC) tissues were mapped to the ginger genome with Hisat2 v.2.0.4 ^105^ alignment program. For *de novo* gene annotation, transcripts well-supported i.e., identified both by the homology-based and the RNA-seq based predictions were selected for *ab initio* prediction using AUGUSTUS v.3.2.3 ^106, 107^. The exon-intron structure of the genes was predicted using, Genscan ^108^ and SNAP ^109^. The results from the three approaches were consolidated using MAKER v.2.31.9 ^110^ to generate a protein-coding gene set. For functional information, *in silico* translated products of coding genes were aligned to seven known protein databases of NR ^111^, InterPro ^112^, GO ^113^, KEGG ^114^, Swissprot and TrEMBL^115^, COG ^116^.

### ncRNA annotation

Four types of ncRNA were annotated in the assembled genome including microRNA (miRNA), transfer RNA (tRNA), ribosomal RNA (rRNA), and small nuclear RNA (snRNA). Non-coding RNAs were annotated based on *de novo*/or homology search methods. The tRNAs genes were annotated using tRNAScan-SE v.1.3.1 ^117^ with default parameters and filtered to remove pseudo annotated tRNA genes. To identify rRNA genes, the *B. rotunda* genome assembly searched against rRNA template sequences (Rfam database, release 13.0) ^118^ of *A. thaliana*, *O. sativa*, and *M. acuminata* with BLASTN with an identity cutoff of >= 90% and a coverage at 80% or more. Using Infernal v.1.1.2 ^119^, mapping of the *B. rotunda* genome sequences to the Rfam database was done to identify miRNA and snRNA genes ^52, 88^.

### Construction of phylogenetic trees

The conserved orthologs genes (COS) in *B. rotunda* genome and 13 other species were identified using Orthofinder program ^61^. Using identified single-copy orthologous genes a neighbour joining (NJ) tree was constructed using MEGAX ^120^ and UpSet plot using UpSetR^121^.

### Gene family expansion and contraction analysis

To identify gene family expansion and contraction, we used the data generated from OrthoFinder as inputs for the Computational Analysis of gene Family Evolution (CAFE)^122^. The phylogenetic tree from OrthoFinder was converted to an ultrametric tree using make_ultrametric.py in OrthoFinder. Gene families with large variance (≥ 100 gene copies) were removed using clade_and_size_filter.py in CAFE package. Divergence times in the phylogenetic tree were estimated using PATHd8 ^123^ calibrated using divergence time between *Brachypodium* and *Oryza* (40–45 million years ago) (The International Brachypodium Initiative 2010) ^124^ and *Arabidopsis* and *Oryza* (130–200 million years ago)^125^. CAFE version 5 ^122^ was used to determine the stochastic birth and death processes and for modelling of the gene family evolution. The parameters for CAFE5 is “cafe5 -i orthofinder_gene_families.txt - t orthofinder_ultrametric.tre -p -e”.

### DNA methylation analysis using bisulfite sequencing (BS-seq)

Bisulfite sequencing reads were pre-processed by trimming low quality reads and adapters by Trim-Galore ^126^ tool specific for bisulfite sequencing. After trimming, bisulfite reads were mapped to draft ginger genome with Bismark ^127^ tool by choosing bowtie aligner with options set to best, minimum map length of 50 bp and insert size of 500bp. Mapping duplicates were removed by Methpipe ^128^ tool. Methcounts program from Methpipe was used for mapping of methylated and unmethylated cytosines where the methylation level at single base resolution was calculated based on the number of 5-methylated cytosines (5mC) in reads, divided by the sum of the C and thymines (T) in CG, CHG and CHH sequence contexts within the coding sequences of all selected genes from *B. rotunda*.

### *De novo* transcriptome assembly of *B. rotunda* and functional annotation

To gather information related to secondary metabolites, expression of genes involved in flavonoid and phenylpropanoid pathways of *B. rotunda* was based on deep transcriptome sequencing of three cell culture types, *in vitro* and *ex vitro* leaves of *B. rotunda*. Based on our previous studies of embryogenesis related genes ^83, 84^ and on levels of metabolites in cell cultures ^42^, we generated deep transcriptome data from five tissue types (each three replicates) to investigate gene regulation patterns in the phenylpropanoid and flavonoid pathways to identify metabolite producing cells in *B. rotunda in vitro* cultured cells. RNA-seq reads were pre-processed using FastQC software (https://www.bioinformatics.babraham.ac.uk/projects/fastqc/) to remove poor quality reads and adapter sequences. The remaining high-quality reads were assembled into contigs using four commonly used short-read assemblers: Oases ^129^; TransABySS ^130^; SOAPdenovo-Trans ^131^ and Trinity ^132^. Two different approaches were used to assemble the transcriptome. In the first approach (best k-mer strategy) high-quality reads were assembled at different k-mer length 21–51 using Oases, TransABySS and SOAPdenovo-Trans whereas the assembly by Trinity used default parameters (K-mer 25). The assemblies from each software in the first approach were further used in the second approach (additive k-mer followed by TGICL) in order to improve the transcriptome assemblies. A two-step strategy was employed for assembly in the second approach in which the contigs generated from all the k-mers by each respective assembler were merged and redundancy was removed using CD-HIT ^133^. The remaining non-redundant contigs were assembled using TGICL clustering tool ^134^ with a maximum identity of 90 and a minimum overlap length of 40. The completeness of the transcriptome assemblies was measured using the BUSCO ^60^ software.

High-throughput functional annotation was performed with Blast2GO Command Line ^135^. To obtain a list of potential homologous for each input sequence, BLAST algorithm (BLASTX) was performed. Blast2GO then maps Gene Ontology (GO) terms associated with the obtained BLAST hits and returns an evaluated functional annotation for the query sequences ^136^. GO mapping and Enzyme Commission (EC) classification were done based on annotation Cut-off 55, E-Value-Hit-Filter 1×10^e-6^, GO Weight of 5, and HSP-Hit Coverage Cut-off 0. The functional enrichment categories among the differentially expressed genes (DEG) were identified by a Fisher exact test with false discovery rate (FDR) cut-off of 0.05. Classification of the *B. rotunda* transcripts into functional categories was performed using the Eukaryotic Orthologous Groups (KOG) ^116^ protein database. *B. rotunda* transcripts were mapped to their biological pathways using the Kyoto Encyclopedia of Genes and Genomes (KEGG) database^137^. Unigenes potentially related to Panduratin A and other secondary metabolites biosynthesis were identified as those with a unigene annotated function matching to enzymes assigned to the flavonoid and phenylpropanoid biosynthetic pathways in the KEGG pathway database.

### Estimation of transcript abundance and differential expression

RSEM software package ^138^ was used for the estimation of the gene expression level with mean fragment length of 200 bp and fragment length standard deviation of 80 bp. The FPKMs (fragments per feature kilobase per million reads mapped) were used to normalize the expression level for each gene and comparison between samples. Bioconductor tool (EdgeR) ^139^ was used for differential expression analysis with a *P*-value threshold of ≤ 0.05 and |log_2_ (Fold Change)| ≥ 1 used to identify significant differential expression of the transcripts.

### Mining of simple sequence repeats from *B. rotunda* transcriptome and genome assembly

Whole genome assembly and assembled transcriptome sequences were searched for SSRs using a modified Liliaceae simple sequence analysis tool (LSAT) pipeline ^140^. Searches were standardized for mining SSRs from mono to 20 bp with minimum repeat loci of 12 nucleotides. SSRs were classified based on SSR locus length (Class I>20nt and Class II 12-20nt) and nucleotide base composition of the SSR loci (AT-rich, GC-rich and AT-GC balance). Primer pair sequences were developed for each identified SSR loci using the default permanents of the primer 3 (http://bioinfo.ut.ee/primer3) software ^141^. Redundant primers pair were eliminated using perl script developed by Biswas et al ^86^. An electronic polymerase chain reaction (ePCR) ^142^ strategy was applied for mapping and estimating the transferability of the designed primers. Primers were mapped on four genomes viz. *Musa acuminata*, *Musa balbisiana, Musa itinerans* and *Ensete ventricosum* those are the most related plant species of the *B. rotunda.* Maximum 2nt mismatch with two gaps was set as a cut off value for ePCR result filter.

### Wet lab validation of the transcriptome SSR (EST-SSR) and genomic SSR (G-SSR) markers

A total 14 (8 EST-SSR and 6 G-SSR) primer pairs were selected based on their *in silico* transferability result to assess their marker potentiality. Three *B. rotunda*, two *Ensete* and three *Musa* species were used to validate selected primer sets. Fresh leaf samples were harvested from the greenhouse grown plants and total genomic DNA was extracted following the CTAB methods. PCR amplifications were carried out for SSR primer validation under the following conditions: 95 °C for 2 min, 35 cycles at 95 °C for 1 min, 60 °C for 1 min, and 72 °C for 1 min, followed by a final elongation at 72 °C for 10 min. Amplified DNA fragments were run on 2% agarose gels in 1 × Tris–Borate-EDTA (TBE) buffer with 80v for 90 min. A 100-bp molecular ladder was used to estimate the amplicon size.

## Acknowledgements

The work was supported by the High Impact Research Chancellery Grant UM.C/625/1/HIR/MOE/SC/15 from the University of Malaya, Kuala Lumpur, Malaysia. We acknowledge our colleagues at the Centre for Research in Biotechnology for Agriculture (CEBAR) and the Plant Biotechnology Research Laboratory (PBRL), University of Malaya and from the Department of Genetics and Genome Biology, University of Leicester, for their kind support and guidance during this research and CEBAR grants TU002G-2018 and RU004A-2020 (for support in research facilities and maintenance).

## Author contributions

Jennifer Ann Harikrishna, Norzulaani Khalid, J. S. (Pat) Heslop-Harrison and Trude Schwarzacher, conceived and designed the study. Sima Taheri, Teo Chee How, Tan Yew Seong, Manosh Kumar Biswas, Naresh V. R. Mutha, Wee Wei Yee and Gan Han Ming, performed the data acquisition, genome sequence assembly and bioinformatics analyses. Trude Schwarzacher and Yusmin Mohd Yusuf performed the chromosome analysis. Sima Taheri, Teo Chee How, Jennifer Ann Harikrishna and J. S. (Pat) Heslop-Harrison wrote the manuscript. All authors assisted with editing of the manuscript and approved the final version.

## Availability of data

Raw sequence data used for genome assembly, mRNA sequencing (RNA-Seq) and whole-genome bisulfite sequencing (BS-Seq) are available at NCBI under BioProject ID PRJNA712941.

