## Supplementary Figures and Tables for "Genome assembly and analysis of the flavonoid and phenylpropanoid biosynthetic pathways in Fingerroot ginger (*Boesenbergia rotunda*)"

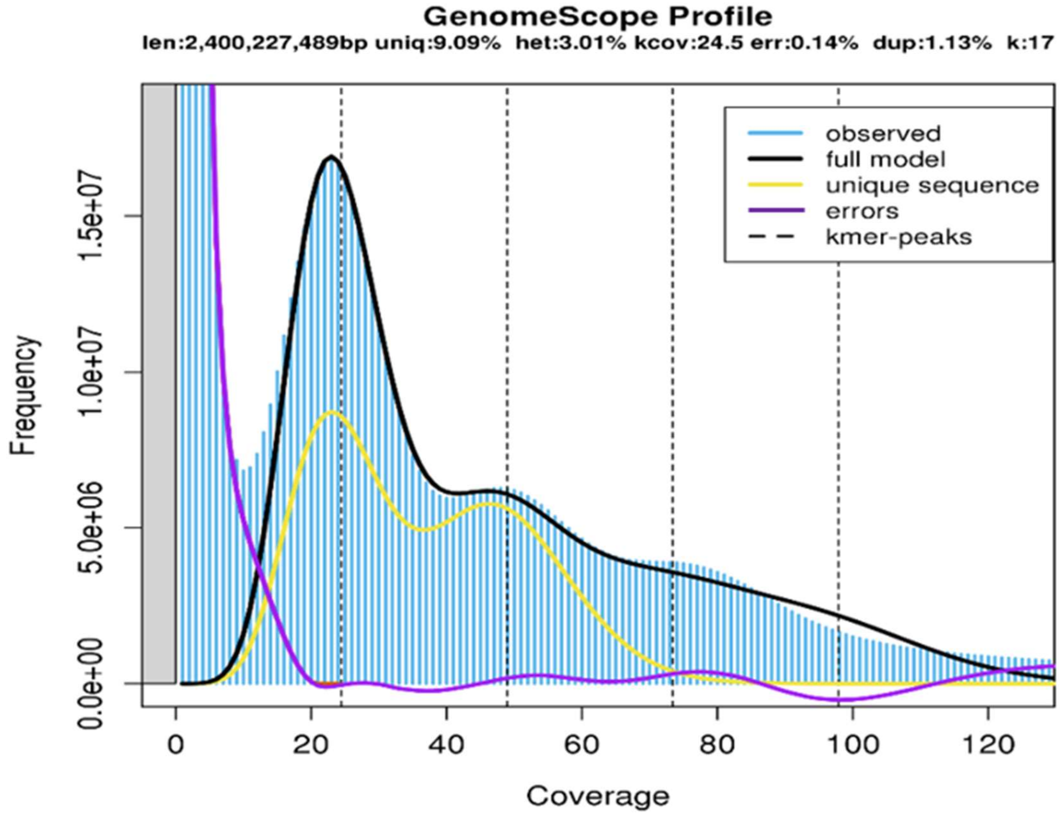


Figure S1. Estimation of *B. rotunda* genome size and heterozygosity of *B. rotunda* genome by GenomeScope 2.0, based on k-mer=17.

*Glycine max* + *B. rotunda*
Modal:
28.90/11.38
= 2.54

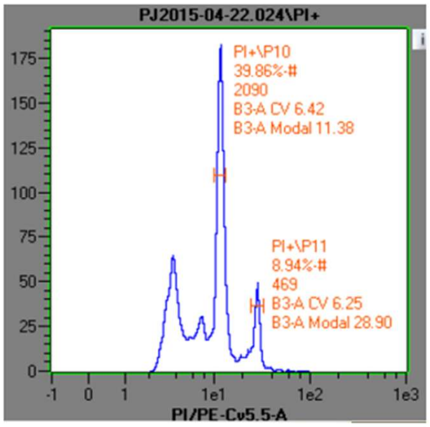


*Zea mays* +
*B. rotunda*
Modal:
51.64/22.49
= 2.30

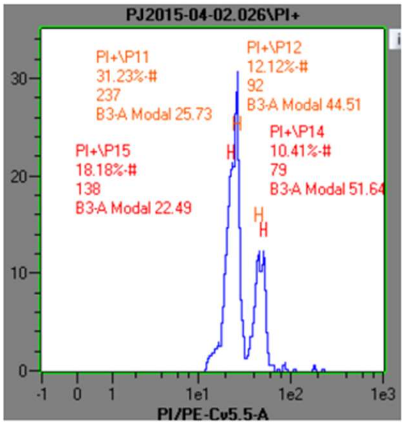


*Pisum sativum* + *B. rotunda* Modal:
45.2/29.81
= 1.52

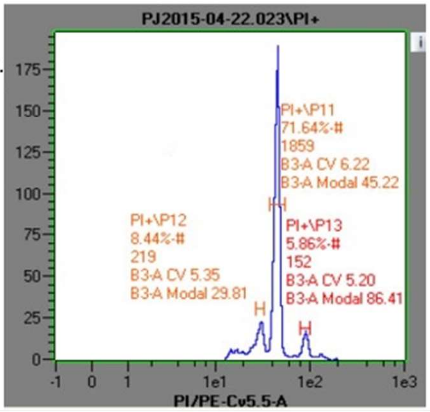


Figure S2. Estimation of absolute nuclear DNA amount (genome size) in *B. rotunda*. The histogram of relative DNA content was obtained after flow cytometric analysis of propidium iodide-stained nuclei of *B. rotunda* and soybean (A), *B. rotunda* and maize (B), and *B. rotunda and pea* (C) which were isolated, stained and analysed simultaneously. i. *Glycine max* cv. Polanka (G) 2C = 2.50 pg DNA, ii. *Zea mays* CE-777 (Z) 2C = 5.43 pg DNA, iii. *Pisum sativum* cv. Ctirad (P) 2C = 9.09 pg DNA were used as internal standards with known genome size.

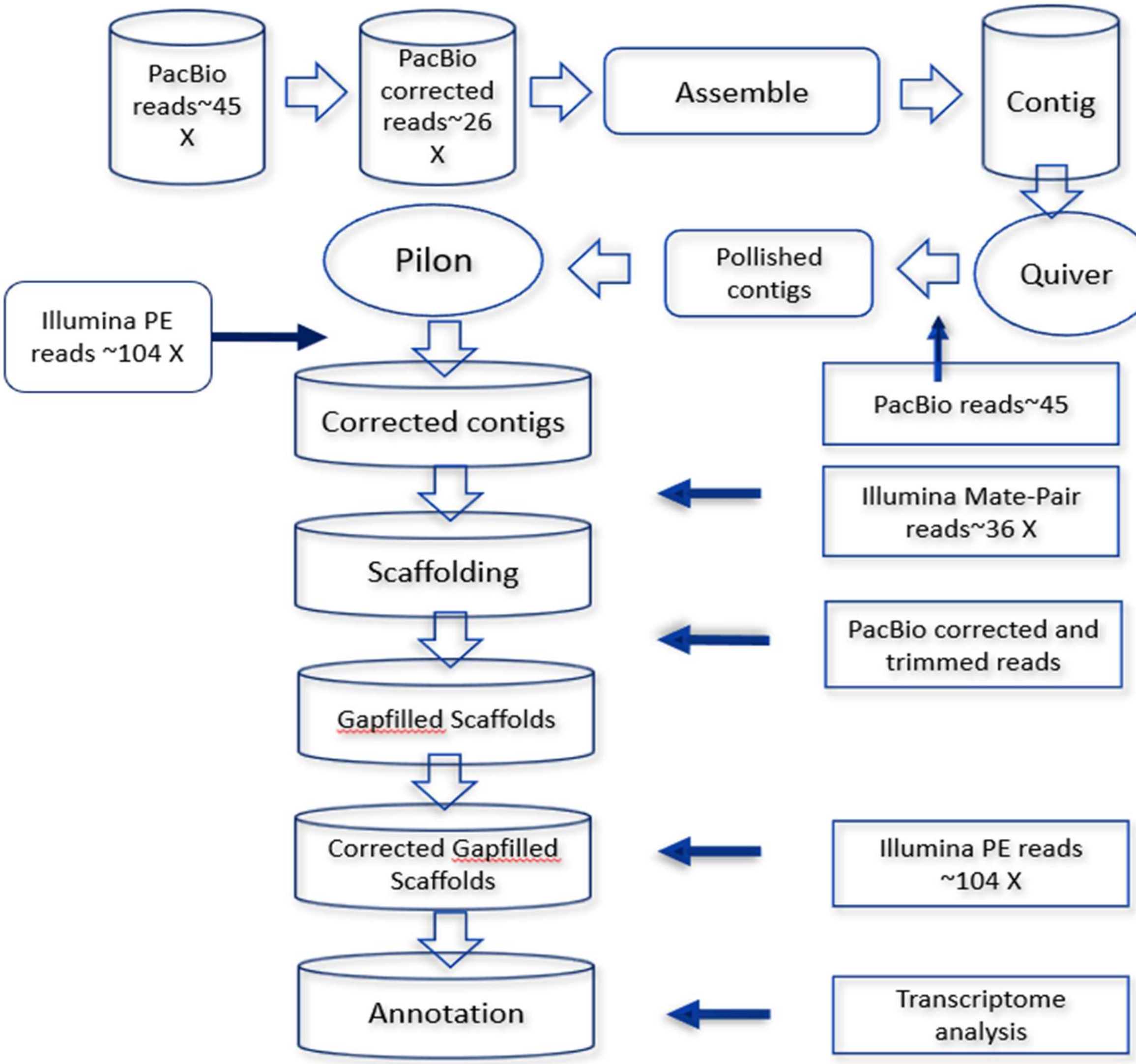


Figure S3. Overview of the processing pipeline used for the assembly of the *B. rotunda* genome

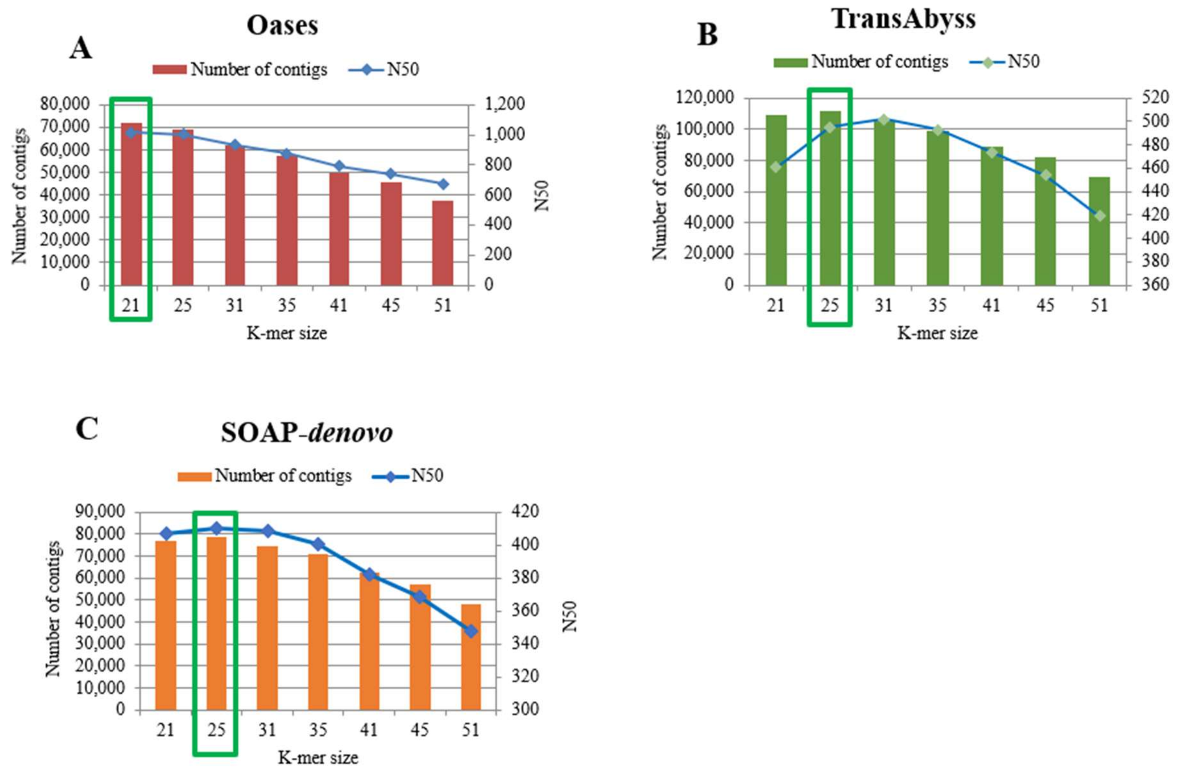


Figure S4. Comparison of total contig number and N50-length obtained after assembly at different K-mer sizes from 21 to 51. Figures A, B, and C represent the contig assembly results of *B. rotunda* from Oases, TransAbyss, and SOAP-*denovo* assemblers. The bars indicate the total number of contigs assembled (Primary axis). The blue line represents the N50 contig length.

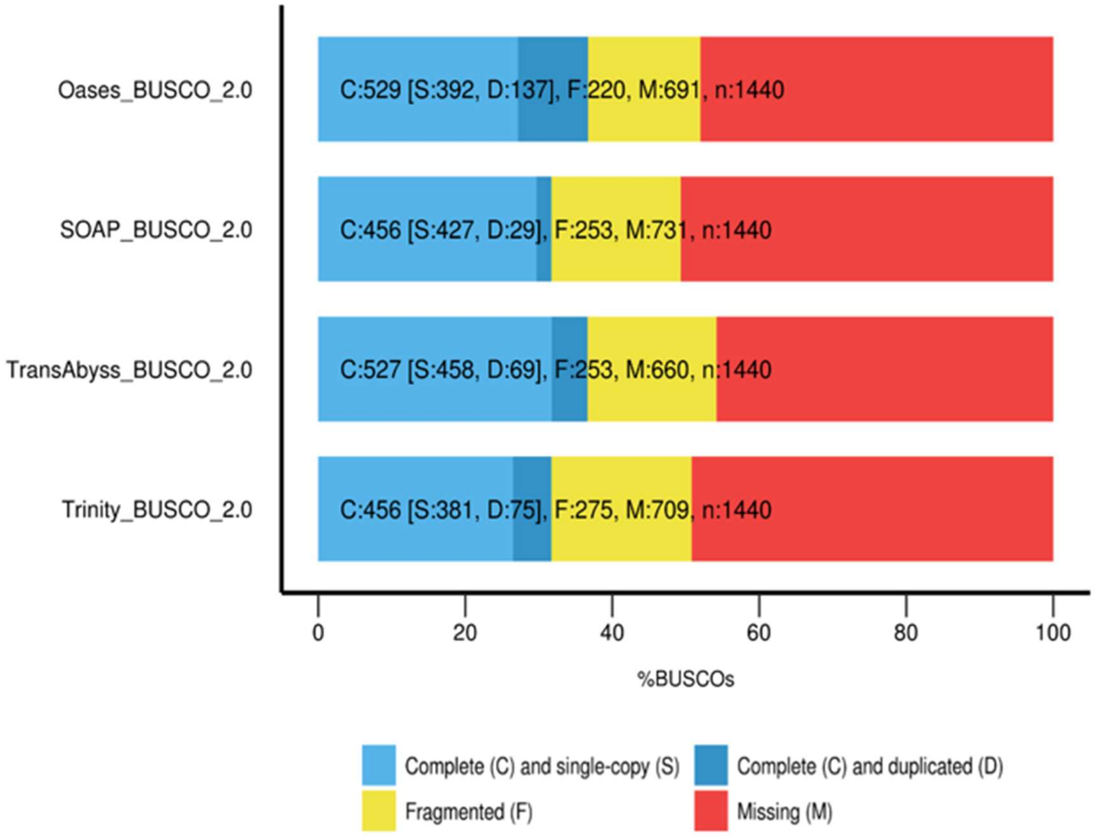


Figure S5. Transcriptome assembly quality assessment of Oases, SOAP-*denovo*, TransAbyss, and Trinity assemblies produced by Benchmarking Universal Single-Copy Orthologs (BUSCO). In each bar, the pale-blue color indicates the number of genes that were detected completely as single-copy genes in the genome, the dark-blue color indicates the number of genes that were detected as single-copy genes but were duplicated, the yellow color indicates the number of genes that were detected as single-copy genes but not completely, and the red color indicates the number of single-copy genes that were not detected among the plant universal single-copy orthologues.

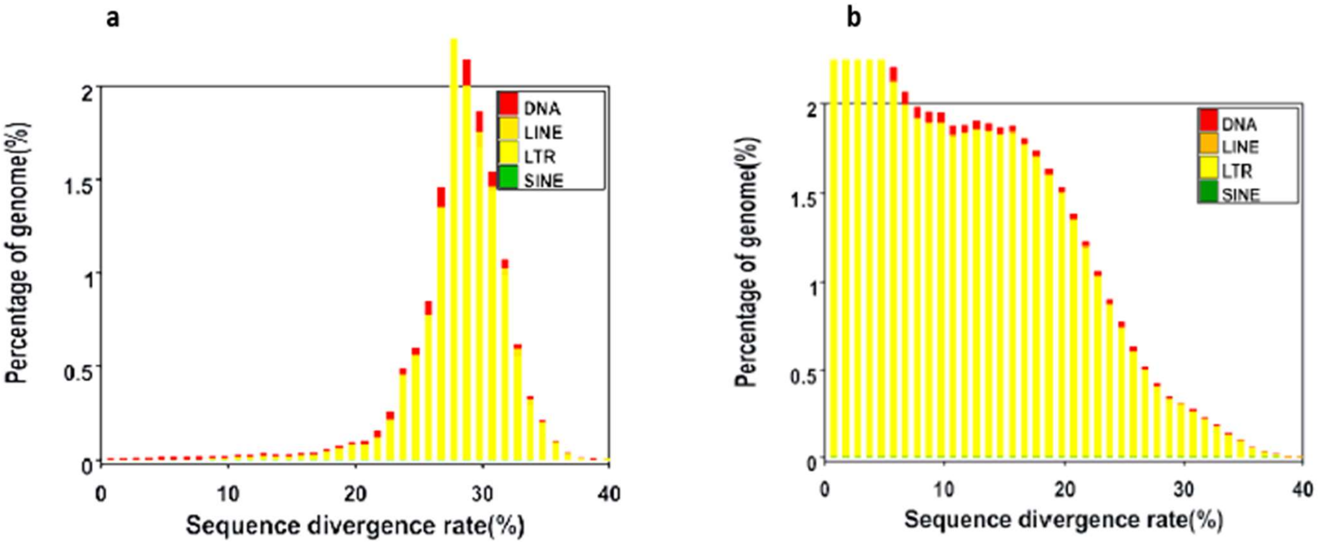


Figure S6. Distribution of divergence rate of each type of TE. The divergence rate was calculated between the identified TE elements in the genome by homology-based method and the consensus sequence in the Repbase (a), and by *de novo* method and the consensus sequence in the predicted TE library (b).

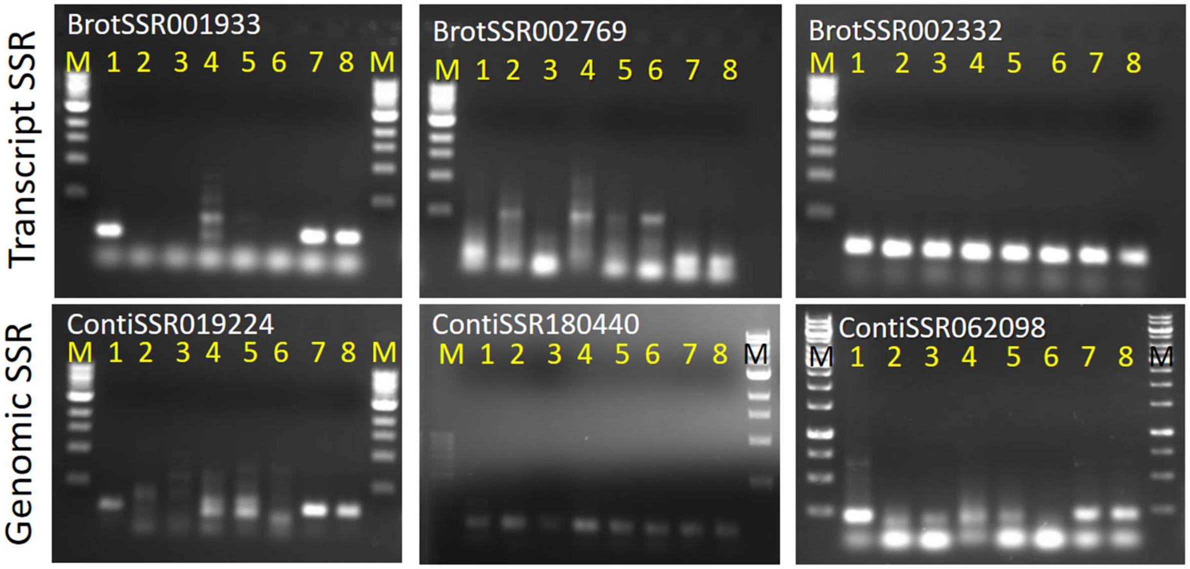


Figure S7. PCR profile of the genomic and transcript derived SSR markers of *B. rotunda*. M: 100bp ladder; 1,7,8: *B. rotunda;* 2, *Musa acuminate, 3. Musa balbisiana, 4. Musa Itinerans* 5,6: *Ensete ventricosum*.

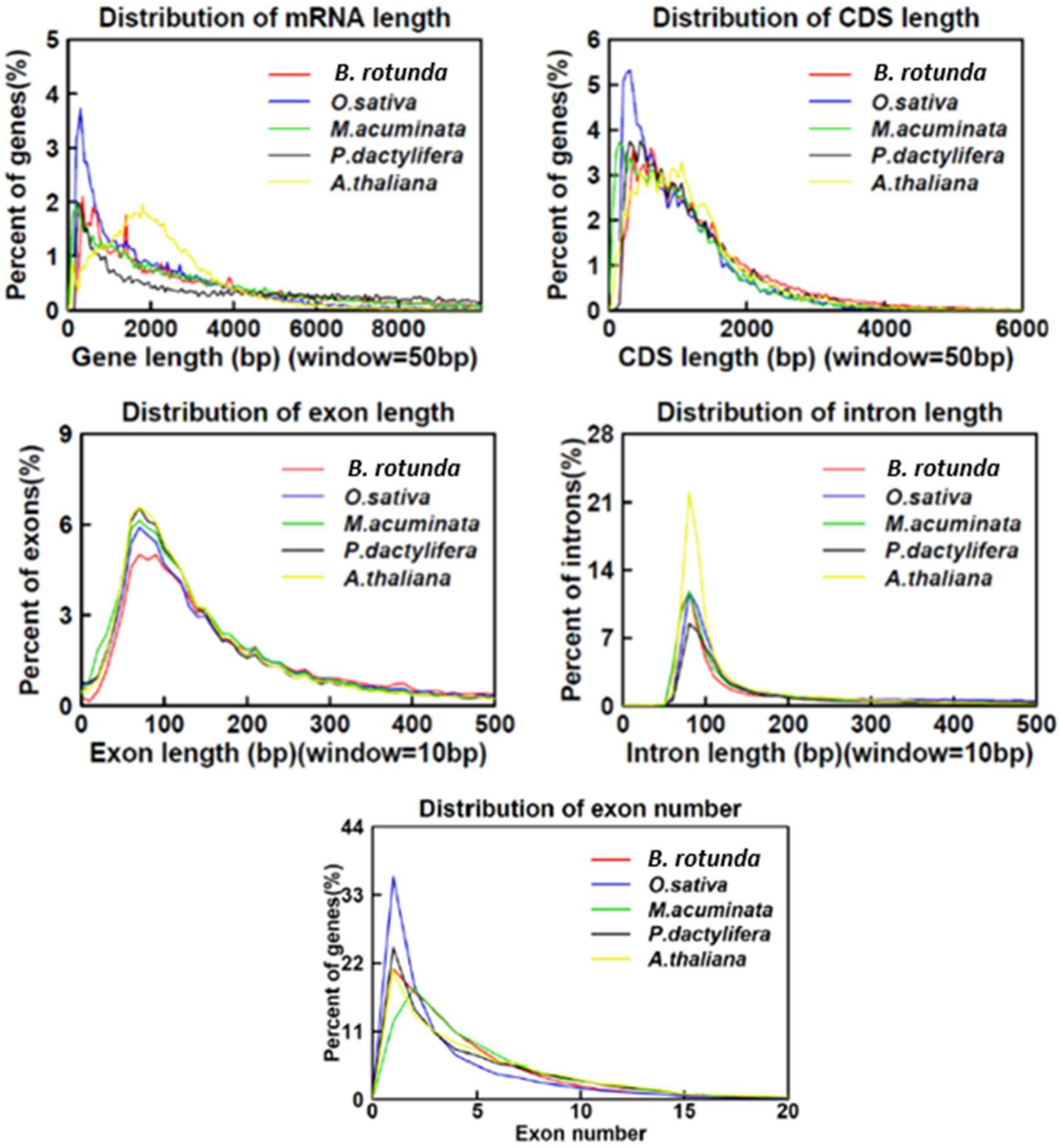


Figure S8. Compared the distribution of several features of the final gene set to homolog species. Window means the length of every point in the horizontal ordinate.

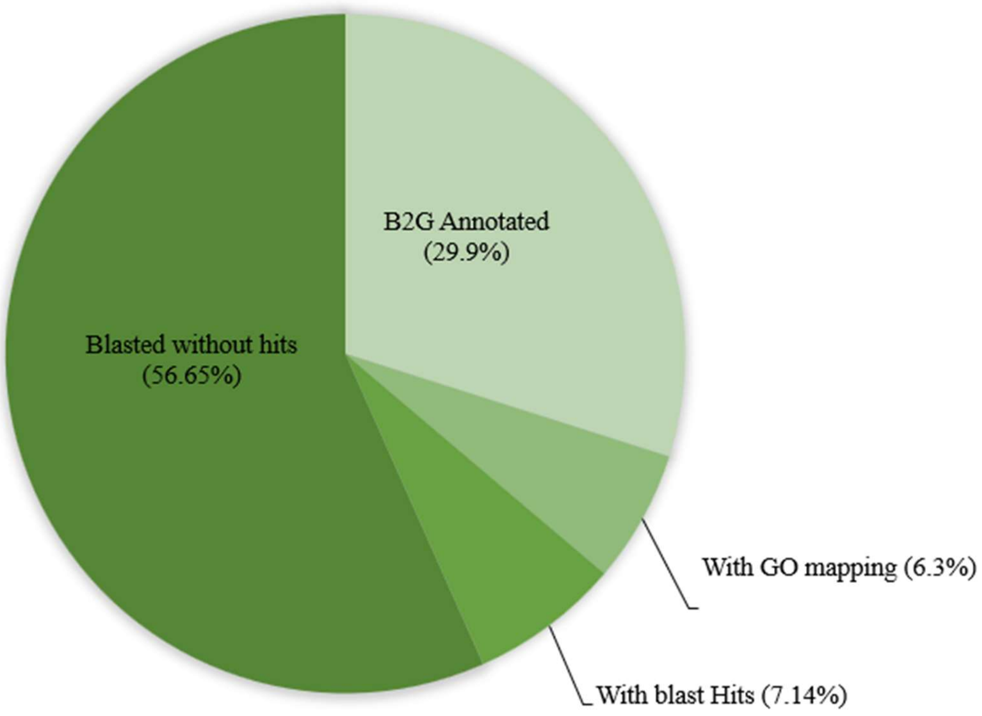


Figure S9. The data distribution pie chart shows the number of sequences which could finally be annotated in comparison to the ones not annotated due to missing results in the blast, mapping, or annotation step.

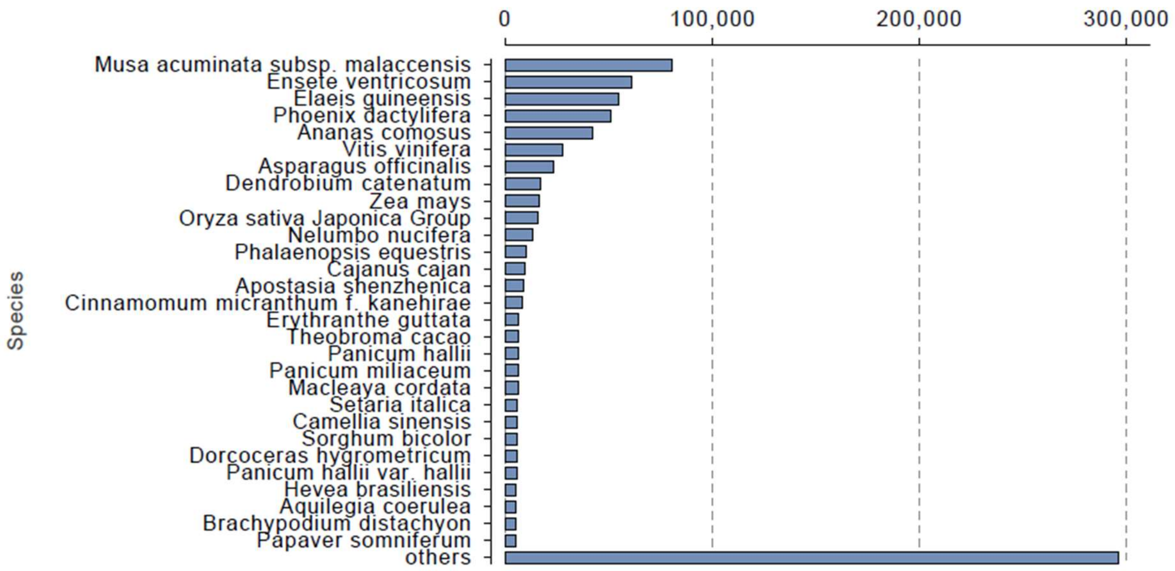


Figure S10. Species distribution of the matched transcriptome sequences *of B. rotunda* in Genebank non-redundant (NR) database. Showing dominant match to *Musa acuminata* subsp.

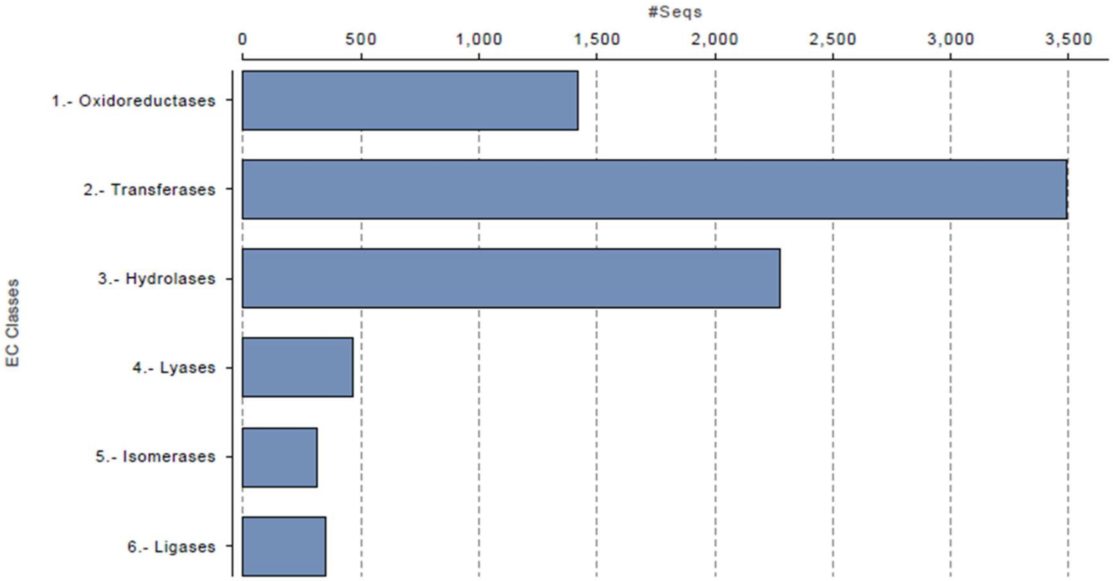


Figure S11. Main Enzyme Code level distribution of *B. rotunda* (Blast2GO)

Table S1. Statistics of Clean data of genome assembly of *B. rotunda*

| Species | Lib ID | Insert size (bp) | Read length (bp) | Data (Gb) | Sequence depth (X) |
|---|---|---|---|---|---|
| *B. rotunda* | WHAXPI019351-103 | 450 | 250_250 | 146 | 58.40 |
| | WHAXPI019352-106 | 450 | 250_250 | 114 | 45.60 |
| | WHGINhgvRAAADWAAPEI-108 | 2000 | 49_49 | 15.5 | 6.20 |
| | WHGINhgvRAABDWAAPEI-109 | 2000 | 49_49 | 15 | 6.00 |
| | BOEnylDAADLAAPE | 5000 | 49_49 | 24.7 | 9.88 |
| | BOEnylDAADTAAPE | 10000 | 49_49 | 13.9 | 5.56 |
| | WHGINebkRAAADUABPEI-101 | 20000 | 49_49 | 2.70 | 1.08 |
| | WHGINebkRAABDUABPEI-102 | 20000 | 49_49 | 6.90 | 2.76 |
| | WHGINhgvRBAADVABPEI-21 | 40000 | 49_49 | 11.40 | 4.56 |
| Total | | | | 350.10 | |

Table S2. Evaluation of completeness of the final assembly by BUSCO

| Summary | C:85.6%[S:60.8%, D:24.8%],F:1.8%,M:12.6%,n:1440 |
|---|---|
| Complete BUSCOs (C) | 1232 |
| Complete and single-copy BUSCOs (S) | 875 |
| Complete and duplicated BUSCOs (D) | 357 |
| Fragmented BUSCOs (F) | 26 |
| Missing BUSCOs (M) | 182 |
| Total BUSCO groups searched | 1440 |

Table S3. List of the primer used for wet-lab validation

| Primer ID | F (5'--3') | R (5'--3') | PCR product size | Tem | Source |
|---|---|---|---|---|---|
| BrotSSR002332 | TGCGTATTGGTACTTATCGGG | ACACGATACCAAGGCAAACC | 243 | 60 | EST |
| BrotSSR001220 | GCTCGATCTCTCGGTACTGG | GTCGCTAGGAATGGATCAGC | 188 | 60 | EST |
| BrotSSR001310 | TAGAAGTCGGCGGAGAAGAA | CTGCTTGAGCACCTTGAAGA | 199 | 60 | EST |
| BrotSSR001518 | AGTCGACCACCATCTCCAAG | CGGAGTTAAGAGCCTCGTTG | 228 | 60 | EST |
| BrotSSR001628 | ATGAACCAGCCAGCTTATGG | CTGGACTGGTCCTTGGAGAG | 122 | 60 | EST |
| BrotSSR001933 | GTTGATCGGCTGGATCACTT | GAAGAGGCTTTGCCTCACTG | 184 | 60 | EST |
| BrotSSR002607 | ATCGTCATCTTCGTCGTCCT | CTCCTCGAATCGCCTCAA | 208 | 60 | EST |
| BrotSSR002769 | GAAGGTGAAGGAGAAGGGCT | CCTCCCCTGTTTCTTCTTCC | 155 | 60 | EST |
| ContiSSR062098 | CGTGAGGATGCTAGAGGAGG | TCTTCAGCCCTTCGATCACT | 220 | 60 | Genomic |
| ContiSSR052791 | CTGTTCACCATCTCCCACCT | CAAGGAGCAAGAATTCGGAG | 142 | 60 | Genomic |
| ContiSSR169255 | CAGCTGATGAGGGATACGGT | TGCCTGCCTTTCTTTGAACT | 217 | 60 | Genomic |
| ContiSSR024743 | ATCTTCCTCATCGTCGTCGT | AGAGAGAGGGAGATGGAGGC | 201 | 60 | Genomic |
| ContiSSR180440 | ATATGGAATGAACACCCCCA | GCTGCGCTTAAATTCCATTC | 160 | 60 | Genomic |
| ContiSSR019224 | CATCGTCCTCTTCTTCCTCG | TGATCCCTCATCGCTCTCTT | 268 | 60 | Genomic |

Table S4. General statistics of predicted protein-coding genes

| Gene set | | Number | Average transcript length (bp) | Average CDS length (bp) | Average exon per gene | Average exon length (bp) | Average intron length (bp) |
|---|---|---|---|---|---|---|---|
| De novo | AUGUSTUS | 185,254 | 3,325 | 892 | 3.28 | 272 | 1,068 |
| | GENSCAN | 133,865 | 3,542 | 453 | 2.76 | 164 | 1,753 |
| | SNAP | 352,601 | 1,934 | 483 | 2.66 | 181 | 872 |
| Homolog | *M.acuminata* | 80,748 | 2,335 | 754 | 3.21 | 235 | 716 |
| | *P.dactylifera* | 165,212 | 1,775 | 745 | 2.24 | 333 | 832 |
| | *O.sativa* | 108,583 | 1,766 | 725 | 2.38 | 304 | 753 |
| | *A.thaliana* | 83,632 | 1,933 | 721 | 2.71 | 266 | 709 |
| RNA-Seq | | 36,033 | 6,773 | 2116 | 7.69 | 275 | 696 |
| MAKER | | 73,102 | 4,312 | 1360 | 4.49 | 303 | 812 |

Note: The average transcript length does not contain UTR. Three approaches were employed in gene prediction: Homolog (*M. acuminata*, *P. dactylifera*, *O. sativa and A. thaliana*), *De novo* (*GENSCAN*, *AUGUSTUS, SNAP*), RNA-seq. Results can be consolidated using the program *maker-2.31.8*.

Table S5. Details of ortholog genes of B. rotunda and other 13 species

| | |
|---|---|
| Number of species | 13 |
| Number of genes | 1007916 |
| Number of genes in orthogroups | 930548 |
| Number of unassigned genes | 77368 |
| Percentage of genes in orthogroups | 92.3 |
| Percentage of unassigned genes | 7.7 |
| Number of orthogroups | 62101 |
| Number of species-specific orthogroups | 29705 |
| Number of genes in species-specific orthogroups | 159786 |
| Percentage of genes in species-specific orthogroups | 15.9 |
| Mean orthogroup size | 15 |
| Median orthogroup size | 5 |
| G50 (assigned genes) | 42 |
| G50 (all genes) | 38 |
| O50 (assigned genes) | 6094 |
| O50 (all genes) | 7052 |
| Number of orthogroups with all species present | 7276 |
| Number of single-copy orthogroups | 0 |

Table S6. Potential unigenes related to biosynthesis of panduratin A and other metabolites in the flavonoid pathway. Samples; *ex vitro* leaf (EVL) = A, *in vitro* leaf (IVL) = B, embryogenic callus (EC) = C, dry callus (DC) = D, watery callus (WC) = E. Note: NDE; Not differentially expressed.

| | **Enzyme name** | **EC number** | **Enzyme class** | **Total Unigene** | **A vs B** | | **A vs C** | | **A vs D** | | **A vs E** | |
|---|---|---|---|---|---|---|---|---|---|---|---|---|
| | | | | | **A** | **B** | **A** | **C** | **A** | **D** | **A** | **E** |
| CL4748Contig1 | Chalcone synthase (CHS) | 2.3.1.74 | Transferase | 1 | NDE | NDE | Down | UP | Down | UP | Down | UP |
| Locus_19369_Transcript_1/1_Confidence_1.000_Length_328 | Chalcone isomerase (CHI) | 5.5.1.6 | Isomerase | 1 | NDE | NDE | Down | Up | NDE | NDE | NDE | NDE |
| CL2047Contig1 | Dihydroflavonol 4-reductase (DFR) | 1.1.1.219 | Oxidoreductase | 1 | Up | Down | Up | Down | Down | Up | Down | Up |
| Locus_7239_Transcript_1/3_Confidence_0.846_Length_1481 | 6′-deoxychalcone synthase (6'DCHS) | 2.3.1.170 | Transferase | 1 | Down | Up | NDE | NDE | NDE | NDE | UP | Down |
| CL15839Contig1 | Hydroxycinnamoyl-CoA shikimate/quinate hydroxycinnamoyltransferase (HCT) | 2.3.1.133 | Transferase | 1 | Down | Up | NDE | NDE | Down | Up | Down | Up |
| CL14369Contig1 | flavanone-3-hydroxylase (F3H) | 1.14.11.9 | Oxidoreductase | 1 | Up | Down | Up | Down | NDE | NDE | NDE | NDE |
| Locus_5053_Transcript_2/5_Confidence_0.462_Length_769 | caffeoyl-CoA O-methyltransferase (CCOAOMT) | 2.1.1.104 | Transferase | 1 | Down | UP | Down | UP | Down | UP | Down | UP |
| CL2457Contig1 | Cinnamyl-alcohol dehydrogenase (CAD) | 1.1.1.195 | Oxidoreductase | 1 | NDE | NDE | Down | Up | Down | Up | Down | Up |
| CL7036Contig1 | 4-coumarate–CoA ligase (4CL) | 6.2.1.12 | Ligase | 1 | Down | Up | NDE | NDE | NDE | NDE | NDE | NDE |
| Locus_2904_Transcript_6/10_Confidence_0.630_Length_2405 (LPO1) | | | | | Up | Down | Up | Down | Up | Down | Up | Down |
| CL2460Contig1 (LPO2) | | | | | Up | Down | Up | Down | Up | Down | Up | Down |
| CL15961Contig1 (LPO3) | | | | | Up | Down | Up | Down | Up | Down | Up | Down |
| CL7193Contig1 (LPO4) | | | | | Down | Up | Up | Down | Down | Up | Up | Down |
| Locus_39965_Transcript_1/1_Confidence_1.000_Length_689 (LOP5) | Lactoperoxidase (LPO) | 1.11.1.7 | Oxidoreductase | 151 | Down | Up | NDE | NDE | Down | Up | Down | Up |
| CL7028Contig1 (LPO6) | | | | | Up | Down | Up | Down | Up | Down | Up | Down |
| Locus_72085_Transcript_1/1_Confidence_1.000_Length_548 (LPO7) | | | | | NDE | NDE | NDE | NDE | Up | Down | Up | Down |
| CL16168Contig1 (LPO8) | | | | | NDE | NDE | NDE | NDE | NDE | NDE | NDE | NDE |
| Locus_2353_Transcript_1/2_Confidence_0.750_Length_911 (LPO9) | | | | | NDE | NDE | NDE | NDE | NDE | NDE | NDE | NDE |
| CL298Contig1 | Phenylalanine ammonia-lyase (PAL) | 4.3.1.25/4.3.1.24 | Lyase | 1 | Down | Up | Down | Up | Down | Up | Down | Up |
| Locus_4538_Transcript_2/2_Confidence_0.667_Length_1890 | coniferyl-aldehyde dehydrogenase (CALDH) | 1.2.1.68 | Oxidoreductase | 1 | Up | Down | Up | Down | Up | Down | Up | Down |
| CL204Contig2 | beta-glucosidase (BGLU) | 3.2.1.21 | Hydrolase | 1 | NDE | NDE | Down | Up | Down | Up | Down | Up |
| Locus_2577_Transcript_6/7_Confidence_0.500_Length_1307 | caffeic acid 3-O-methyltransferase (COMT) | 2.1.1.68 | Transferase | 1 | Up | Down | Up | Down | NDE | NDE | Up | Down |
| CL646Contig1 | Cinnamoyl-CoA reductase (CCR) | 1.2.1.44 | Oxidoreductase | 1 | NDE | NDE | Up | Down | NDE | NDE | NDE | NDE |
